# How helpful are the protein-protein interaction databases and which ones?

**DOI:** 10.1101/566372

**Authors:** Akhilesh Kumar Bajpai, Sravanthi Davuluri, Kriti Tiwary, Sithalechumi Narayanan, Sailaja Oguru, Kavyashree Basavaraju, Deena Dayalan, Kavitha Thirumurugan, Kshitish K Acharya

**Affiliations:** Structural Biology Lab, Centre for Biomedical Research, School of Bio Sciences & Technology (SBST), Vellore Institute of Technology (VIT) University, Vellore - 632014, Tamil Nadu, India; Shodhaka Life Sciences Pvt. Ltd., Electronic city, Phase I, Bengaluru (Bangalore) - 560100, Karnataka, India (www.shodhaka.com); Biological data Analyzers’ Association (BdataA), Electronic City, Phase I, Bengaluru (Bangalore) - 560100, Karnataka, India (www.startbioinfo.com/BdataA); Insitute of Bioinformatics and Applied Biotechnology (IBAB), Phase I, Electronic City, Bengaluru (Bangalore) - 560 100, Karnataka, India (www.ibab.ac.in)

**Keywords:** Protein-protein interaction, protein interaction databases, interaction databases, database comparisons, protein interactions, molecular networks, systems biology

## Abstract

Protein-protein interactions (PPIs) are critical, and so are the databases and tools (resources) concerning PPIs. But in absence of systematic comparisons, biologists/bioinformaticians may be forced to make a subjective selection among such protein interaction databases and tools. In fact, a comprehensive list of such bioinformatics resources has not been reported so far. For the first time, we compiled 375 PPI resources, short-listed and performed preliminary comparison of 125 important ones (both lists available publicly at startbioinfo.com), and then systematically compared human PPIs from 16 carefully-selected databases. General features have been first compared in detail. The coverage of ‘experimentally verified’ vs. all PPIs, as well as those significant in case of disease-associated and other types of genes among the chosen databases has been compared quantitatively. This has been done in two ways: outputs manually obtained using web-interfaces, and all interactions downloaded from the databases. For the first approach, PPIs obtained in response to gene queries using the web interfaces were compared. As a query set, 108 genes associated with different tissues (specific to kidney, testis, and uterus, and ubiquitous) or diseases (breast cancer, lung cancer, Alzheimer’s, cystic fibrosis, diabetes, and cardiomyopathy) were chosen. PPI-coverage for well-studied genes was also compared with that of less-studied ones. For the second approach, the back-end-data from the databases was downloaded and compared. Based on the results, we recommend the use of STRING and UniHI for retrieving the majority of ‘experimentally verified’ protein interactions, and hPRINT and STRING for obtaining maximum number of ‘total’ (experimentally verified as well as predicted) PPIs. The analysis of experimentally verified PPIs found exclusively in each database revealed that STRING contributed about 71% of exclusive hits. Overall, hPRINT, STRING and IID together retrieved ~94% of ‘total’ protein interactions available in the databases. The coverage of certain databases was skewed for some gene-types. The results also indicate that the database usage frequency may not correlate with their advantages, thereby justifying the need for more frequent studies of this nature.

## Introduction

The flood of databases and tools can enforce subjective selections among biologists. Periodic comparative studies are needed to counter this problem. But such studies are not always a priority among biologists or bioinformaticians. Protein-protein interaction (PPI) databases and tools (resources) are not an exception to this. PPIs are important in deciphering the underlying biological mechanisms in various normal and disease conditions [1–2]. Understandably, there has been a substantial increase in the efforts in characterizing them using various experimental techniques [3] and prediction methods [4]. The time line data in PubMed indeed shows a constant increase in the number of research articles on PPIs across years. With increase in data, many scientists realized the need to compile the PPIs, and there seem to be a large number of PPI databases and computational tools (henceforth collectively referred as ‘resources’) that enable storing, accessing, and analyzing the interaction data in various contexts. The databases can be identified into three broad types based on the type of interaction data collected: i) experimentally verified (EV) PPIs, either curated from literature or submitted directly by the authors, ii) PPIs predicted through various computational methods, and iii) both curated and predicted PPIs. The databases that independently compile the data are called as primary databases, and the ones that collect data from multiple primary databases are called as meta-databases.

Scientists are also increasingly using PPI resources for various purposes – particularly with the advent of genome-wide study methods for biomolecules. Bioinformaticians have also been putting efforts to make a better use of primary databases in terms of improved predictions, visualizations, network analysis etc. While it is a boon for the scientific community to have many specialized protein interaction databases and software, objectively selecting a specific resource is often a daunting task for the biologists. This is most probably true for most bioinformatics resources. Most of the time, researchers may have to make an arbitrary choice by selecting one of the databases or tools used by their peers, as the usage of such bioinformatics resources is usually one of the small component in their research work and they cannot afford to deviate from their main research goals. It is also important to note that the databases and tools are too many to compare (see www.startbioinfo.com). In fact, the actual number of PPI-resources had not been reported anywhere till now, even though there have been some efforts to compile (e.g., www.startbioinfo.com and http://pathguide.org/) or compare them. Tuncbag et al., [5] have compiled the features of about 80 PPI databases, protein interface databases and tools, binding site prediction servers and protein docking tools and servers. The authors have also shown the use of protein interaction resources using specific case studies. Further, Zhou and He have described about 25 tools that retrieve protein-protein interactions from literature, through text mining [6]. About 50 PPI resources have been classified in Startbioinfo (http://startbioinfo.com/cgi-bin/resources.pl?tn=PPI CUS) based on their applications/utilities.

There have also been good efforts to quantitatively compare the performance and/or capacities of the PPI databases. Turinsky et al., [7] have quantified the agreement between curated interactions shared across nine major public databases (BIND, BioGRID, CORUM, DIP, HPRD, IntAct, MINT, MPact, and MPPI). Another study [8] analyzed the extent of coverage of interactions across multiple species by comparing six protein interaction databases (BIND, BioGRID, DIP, HPRD, IntAct and MINT). In 2006, Mathivanan et al., [9] compared the primary protein interaction databases containing literature curated PPIs for human proteins (BIND, DIP, HPRD, IntAct, MINT, MIPS, PDZBase and Reactome) and quantified the coverage of PPIs and proteins. They also compared the performances of these databases using 16 genes/proteins. But there is a need for further work.

Overall, while it is important to systematically compare protein-interaction databases, the highest number of databases compared systematically is only 9, reported in 2010 [7]. A general compilation of such resources has been a maximum of 89 and reported in 2009 [5]. While the above-mentioned comparative studies provided useful insights on the coverage of different PPI databases, such studies need to be conducted periodically as some of the resources would be updated over time, a few may stop working and new ones are likely to be published every year. It also should be noted that earlier studies primarily focused on the overall coverage in terms of the content of the PPI databases but missed an important user-perspective, a quantitative comparison of response of the PPI databases to specific types of biological queries via the web-interfaces. Comparison of databases from user’s perspective using gene-names as queries was reported in one of the published comparative studies in 2006, and with a random set of 16 genes [9]. Such studies are important because, irrespective of the method of data compilation, a biologist would query databases with various types of genes/proteins. It would be interesting, for example, to check the relative performance of the databases when queried with well-studied vs. less studied genes. Similarly, no one has tested the possibility of biases in the curation process used by databases for various diseases. There is also a possibility of unintended bias due to literature selection for auto-text-mining or manual data collection, as different databases and search engines have different efficiency and coverage of the type of articles. For example, about 70% of articles related to Chikungunya disease prevalence that could be scanned by Google Scholar were missed by most other scientific literature search engines [10]. Genes specifically transcribed in different tissues may also be suspected to have different amount of protein interaction studies, and one cannot simply rule out a differential coverage of such gene-sets across databases. For example kidney, uterus and testis tissues have different amount of corresponding research papers available from PubMed, and this might influence the PPI data -corresponding to genes specific to these tissues or the ubiquitously expressed genes. It is not easy to imagine the net result of type of research papers used and the mode of data compilation, as well as the methods of storage, query-handling (including identifier conversions) and algorithms, used by different databases. Hence, web-based querying might be the best way to compare the end-results from users’ perspective.

Moreover, earlier comparative studies mainly focused on primary databases storing curated/experimentally verified protein interactions, whereas, the meta-databases, which are being widely used by researchers, were not considered at all. Thus, there is a need to compile a comprehensive list of PPI resources, carefully select and then systematically compare them from user’s perspectives. We initiated an exhaustive compilation of PPI resources and performed a comprehensive comparative study to evaluate the current features and coverage of a few selected databases. After a thorough initial screening, we selected and evaluated sixteen human protein interaction databases by comparing the interactions obtained through web-interface based query of tissue specific and disease associated genes. We also compared the complete protein interaction data downloaded from the databases, to provide a quantitative measure on the exclusive and shared proteins and interactions across them. In addition, the study describes various features, advantages, and disadvantages of the databases considered.

## Methods

A schematic representation of the overall methodology of the comparative analysis is represented in **Fig 1**.

**Fig 1:**
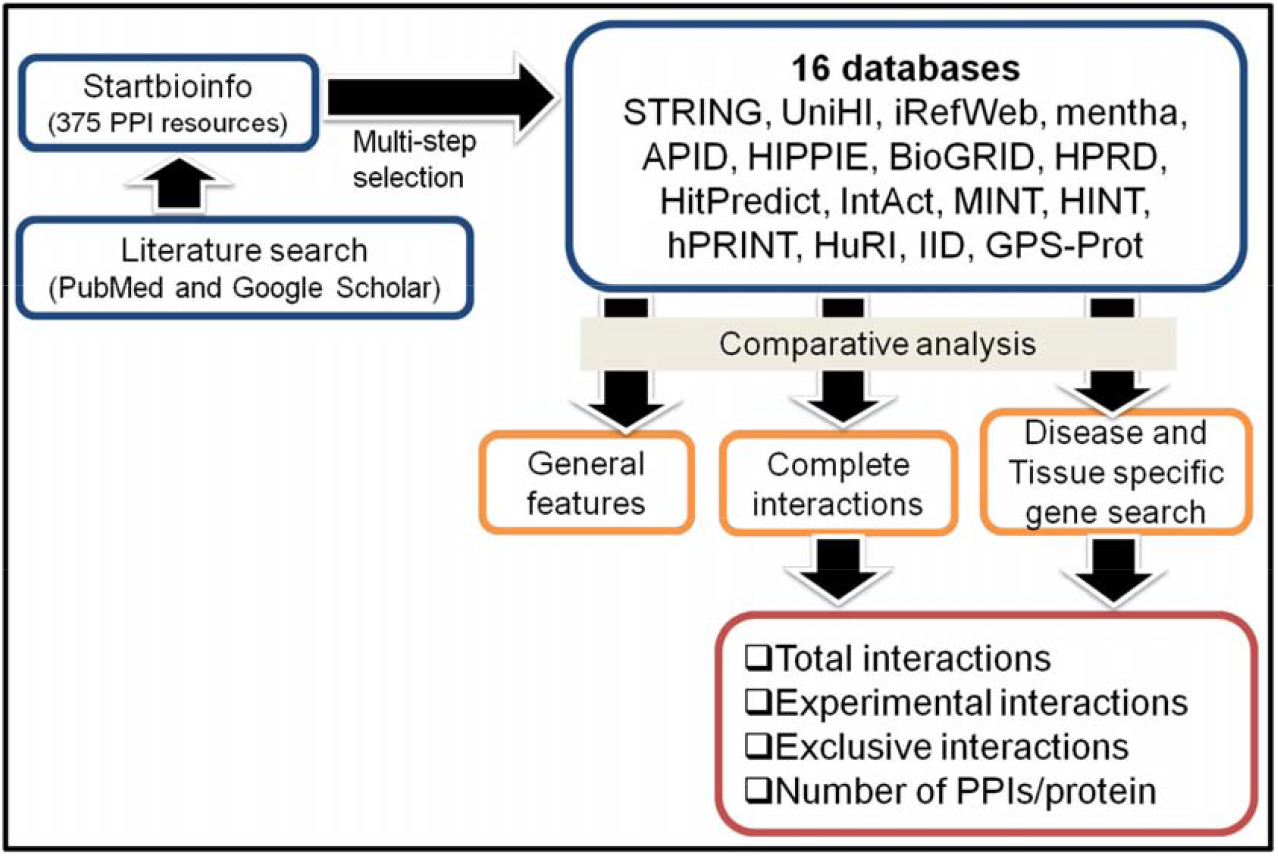
Schematic representation of the methodology used in the comparative analysis of sixteen protein interaction databases.

### Compilation and selection of protein interaction databases

Extensive literature search was performed using PubMed and Google to compile protein interaction resources. Basic features, URL, publication date, and corresponding article(s) for each resource were also listed by browsing through their web-pages. Relative usage frequency was assessed for every resource, and a rank was assigned from 1 to 100. Each rank corresponded to one or more resources with similar usage frequency. Rank 1 was assigned to resources with highest usage whereas 100^th^ rank corresponded to resources that were least commonly used. The usage frequency rank (UFR) was independently calculated by 3 different methods first, ranks were assigned and the average rank was then derived and considered to be the final UFR for each resource. The rankings were also cross-checked by random manual assessment of the citations of the resources, and were confirmed to be indicating a correct relative usage frequency of the resources. Details of the methods used for UFR calculation and rank-assignment are provided in Startbioinfo portal (http://startbioinfo.com/description.html)

The databases used for detailed comparative analysis were selected based on a multi-step selection process: (A) resources having an UFR of 98 and below were shortlisted, (B) the PPI resources that have been considered by the existing comparative studies and (C) the ones published since 2011 were also included in this list, irrespective of their UFR. The resources thus short-listed had a non-redundant set of 125 PPI databases/tools. Each of the 125 PPI resource was then queried with a gene/protein identifier for testing its functionality, and the resources belonging to the following types were excluded from further consideration for the detailed analysis: (i) those which were not functional (35 resources), (ii) protein interaction related tool/software/text-mining systems (32 tools; only PPI databases were retained), (iii) databases that did not provide human PPI data (18 databases), (iv) databases providing only visualization of the protein interaction network or those restricted to very specific type of PPI information, such as compartment specific (e.g., ComPPI), specific to extracellular matrix proteins (e.g., MatrixDB), or specific to mitochondrial proteins (e.g., MitoInteractome). Finally, we obtained 16 PPI databases that were used for the detailed comparative analysis. More details about each database can be found in **Supplementary data, S1 Text**.

### Selection of the gene-sets for database comparison

We considered tissue specific (kidney, testis, and uterus), ubiquitous, and disease associated (breast cancer, lung cancer, Alzheimer’s, cystic fibrosis, diabetes, and cardiomyopathy) genes for querying the 16 selected PPI databases. Genes transcribed ubiquitously or tissue-specifically were prepared using tissue specific gene expression databases for testis [11], uterus [12], and kidney (http://resource.ibab.ac.in/MGEx-Kdb/), which were prepared by manually compiling gene expression data. These databases also assign a reliability score for expression in each tissue and condition. These scores were useful in hierarchically arranging the genes in each list and select the top ones. Genes reported earlier [13] as well as the TiGER [14] and PaGenBase [15] databases were also used. The ubiquitous and tissue specific genes were categorized into well- and less-studied based on the number of published articles in PubMed. More details on the selection of tissue specific and ubiquitous genes can be found in **Supplementary data, S2** Text. The disease associated genes were obtained from Online Mendelian Inheritance in Man (OMIM) database [16].

### Comparison of protein-interaction data: gene based queries

For every gene-set, the ‘total’ (predicted and experimentally verified) and experimentally verified (‘EV’) interactions were obtained by querying individual gene in each database. Both the query and results were restricted to human proteins. In case of STRING, the proteins were queried in two ways: using the web-interface (STRING-Web) and through Application Programming Interface (STRING-API). The API was used to obtain all the available interactions for the query proteins, as the web-interface output is limited to 500 interactions. The IntAct database on the other hand provides interactions for multiple species in one table when queried through the web-interface, which requires additional processing to obtain the human interactors. The protein interactions for each gene/protein were hence compiled from the complete PPI data downloaded from the website. The number of interactions for IntAct obtained through this process and the web-based queries showed no difference which was verified with at least 10 sample proteins. The output from a few databases had identifiers other than official gene symbols (e.g., APID and iRefWeb provided UniProt identifier or accession number), which were converted using Database for Annotation, Visualization and Integrated Discovery (DAVID) [17], and the partners with successful conversion (~95%) were considered.

To assess the coverage of the databases (henceforth referred to as ‘yield-coverage’), first a union list of non-redundant interactions for each protein was derived by considering the interacting partners from all the 16 databases, and was considered as the maximum possible interactions for that particular protein (hereafter referred as ‘Max-PPIs’). The total number of possible interactions for the complete gene-set was derived by adding the ‘Max-PPIs’ for all the genes in that gene-set (hereafter referred as ‘Max-PPIs-GS’), and was considered as reference for calculating the yield-coverage of each database for the gene-set. The yield-coverage of each database was calculated independently for ‘total’ and ‘EV’ interactions by using the following equation:

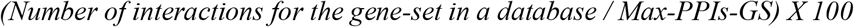

Further, the interactions obtained for each protein were compared across all the databases to obtain the number of exclusive interactions contributed by each database. The total number of exclusive interactions contributed by each database was derived by adding the exclusive interactions for all the proteins in a particular gene-set. Additionally, we calculated the exclusive interactions for each database separately by considering the output of STRING-Web for selected gene-sets (ubiquitous, kidney, breast cancer, Alzheimer’s, and cardiomyopathy). The yield-coverage of each database for exclusive contribution was calculated independently for ‘total’ and ‘EV’ interactions by using the following equation:

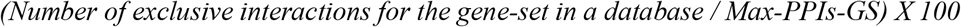

Similarly, the yield-coverage was calculated for the well and less-studied genes by separately considering the interactions for the corresponding genes. Well and less-studied genes were categorized based on the number of articles retrieved.

### Comparison of protein interaction data: complete downloaded data

Complete interaction data from each of the 16 databases except for GPS-Prot (no download link was available) was downloaded during Aug-Sep 2017. In a few cases the complete data downloaded from the database needed some modifications to be considered for the comparison with other databases. In case of HitPredict, the downloaded file did not contain interactions from HPRD, although the output contains data from this resource when a web-based query is used. Hence, the interaction data downloaded from HPRD was included to the data obtained from HitPredict. In case of HuRI database, the ‘preliminary’, or ‘unpublished’ interaction data was not downloaded due to restricted permissions. In case of BioGRID, the genetic interactions were excluded. Further, only binary or physical interactions were considered, wherever applicable. For the databases that did not have a download link for human specific data, complete data was downloaded and the human interactions were extracted based on species name or taxonomy identifier. Cross-species interactions, where one of the interacting proteins was not from human, were removed. The genes/proteins were converted to official gene symbols using Database for Annotation, Visualization and Integrated Discovery (DAVID) [17], wherever applicable. On an average, around 98% of the identifiers were successfully converted to gene symbols (**Supplementary data, Table S1**). Duplicate interaction pairs were removed (interaction pairs such as, P1 – P2 and P2 – P1 were treated same and only one pair was retained) to create a non-redundant list of interactions for each database. The proteins and interactions were then compared across the databases to obtain the number of shared and exclusive contribution. A non-redundant list of interactions from all the databases was considered as the reference for calculating the individual database coverage (henceforth referred to as ‘content-coverage’). The comparison was performed for both ‘total’ and ‘EV’ interactions independently by considering the respective reference interactions. Additionally, the list of proteins corresponding to ‘total’ and ‘EV’ interactions were obtained and compared across the selected databases.

## Results

A total of 375 databases and tools were compiled through extensive literature search and are listed in Startbioinfo web-portal (http://startbioinfo.com/cgi-bin/simpleresources.pl?tn=PPI_AR), along with their basic features. Those with a potential higher relevance to biologists were identified to be 125 and can also be accessed through Startbioinfo portal (http://www.startbioinfo.com/Selected_PPI.xls). We selected 16 protein interaction databases based on a multi-step selection process, for the detailed comparative analysis (**Table 1**). Of the 11 secondary interaction databases considered, 10 integrate data from multiple databases including IntAct and 9 databases integrate data from BioGRID, DIP, and HPRD including others, making these 4 the most preferred primary databases. The databases considered in the current study collect data either by their own curation efforts and depositions of experimentally obtained PPI data by other authors, or by integrating data from other PPI resources. A few databases (e.g., hPRINT, STRING) additionally implement prediction methods to identify interactions. Many of the databases (STRING, mentha, hPRINT, HIPPIE, GPS-Prot, BioGRID, HitPredict, IntAct, MINT, iRefWeb) not only list the interacting proteins but also provide a scale indicating the confidence of interaction, mainly calculated based on number of evidences and the experimental techniques/prediction methods used for identifying the interactions. Very few databases (e.g., STRING, mentha, APID and HuRI) allow the user to import multiple genes/proteins as a query, for which the output can be direct (interactions involving proteins within the list) and/or indirect (interactions involving proteins other than the list) interactions. This would give an opportunity for the user to identify the intermediate proteins that are important for a process or pathway that are not in the query. There are also some unique features provided by some of the databases. For example, STRING is the only database among the 16 that provides information on the mode of action of the interaction. And, IID is the only database that provides information about the expression status (tissue wise) of the interacting proteins. Such information can help in overall understanding of a mechanism where the proteins are known to be interacting, and also aid in further analysis. Among the databases considered, IntAct, MINT, and mentha adhere to the policies of International Molecular Exchange (IMEx) consortium [18]. The IMEx consortium is an international collaboration between a group of major public interaction databases that provides users with a non-redundant set of molecular interactions, collected using a detailed curation model and are available in the Proteomics Standards Initiative Molecular Interactions (PSI-MI) standard formats [19]. A brief overview and features of each of the 16 databases are provided in **Supplementary data, S1** Text. **Fig 2** depicts the data exchange across different primary and secondary databases.

**Table 1:** An overview of protein interaction databases considered for the comparative analysis.

| Database | URL | Version used | Primary/Secondary | Data type (E/P) | Network visualization | Complete data | Organism (number of organisms) | Unique Features |
| --- | --- | --- | --- | --- | --- | --- | --- | --- |
| **STRING [20]** | http://string-db.org | v10.5 | Secondary | E & P | Yes | Yes | Multiple (>10) | Functional enrichment of interacting proteins. Mode of action of the interaction. User-friendly network analysis. |
| **UniHI [21]** | http://www.unihi.org | v7.1 | Secondary | E & P | Yes | Yes<sup>2</sup> | Human<sup>3</sup> | Gene-regulatory interactions. |
| **mentha [22]** | http://mentha.uniroma2.it/ | -- | Secondary | E | Yes | Yes | Multiple (>10) | -- |
| **hPRINT [23]** | http://print-db.org/hprint-web/ | v10.1 | Secondary | E & P | No | Yes | Human | Integration and prediction of PPI through random forest and Bayesian learning approaches. |
| **APID [24]** | http://apid.dep.usal.es/ | vJune 2017 | Secondary | E | Yes | Yes | Multiple (>10) | -- |
| **HIPPIE [25]** | http://cbdm-01.zdv.uni-mainz.de/~mschaefer/hippie/ | v2.1 | Secondary<sup>1</sup> | E | Yes | Yes | Human | Confidence scored and functionally annotated human PPIs. |
| **GPS-Prot [26]** | http://www.gpsprot.org | v3.0.5 | Secondary<sup>1</sup> | E | Yes | No | Human & HIV | Visualization of human and HIV PPIs. |
| **BioGRID [27]** | http://thebiogrid.org | v3.4.154 | Primary | E | Yes | Yes | Multiple (>10) | Genetic interactions |
| **HPRD [28]** | http://www.hprd.org | v9 | Primary | E | No | Yes<sup>2</sup> | Human | -- |
| **HitPredict [29]** | http://hintdb.hgc.jp/http/ | v4 | Secondary | E | Yes | Yes | Multiple (>10) | Experimentally determined physical protein-protein interactions with reliability scores. |
| **IntAct [30]** | http://www.ebi.ac.uk/intact | v4.2.10 | Primary | E | Yes | Yes | Multiple (>10) | Protein chemical interactions. |
| **MINT [31]** | http://mint.bio.uniroma2.it | -- | Primary | E | Yes | Yes | Multiple (>10) | -- |
| **HINT [32]** | http://hint.yulab.org/ | v4 | Secondary | E | Yes | Yes | Multiple (>10) | High quality experimental PPIs. |
| **IID [33]** | http://ophid.utoronto.ca/iid | v2017-04 | Secondary | E & P | No | Yes<sup>2</sup> | Multiple (<10) | Tissue specific annotation of protein interactions |
| **HuRI** | http://interactome.baderlab.org/ | -- | Primary | E | Yes | Yes<sup>2</sup> | Human | High quality mapped PPIs using Y2H and validated using orthogonal assays. |
| **iRefWeb [34]** | http://wodaklab.org/iRefWeb/ | v13.0 | Secondary | E & P | No | Yes | Multiple (>10) |  |

<sup>1</sup>Mainly secondary databases, but also collect interaction data from literature; <sup>2</sup>Registration required for downloading complete or partial data; <sup>3</sup>Multiple species query possible. E: Experimental; P: Predicted. An exhaustive list of protein interaction resources (tools and databases) can be found in **Startbioinfo** (http://startbioinfo.com/cgi-bin/simpleresources.pl?tn=PPI_AR).

**Fig 2:**
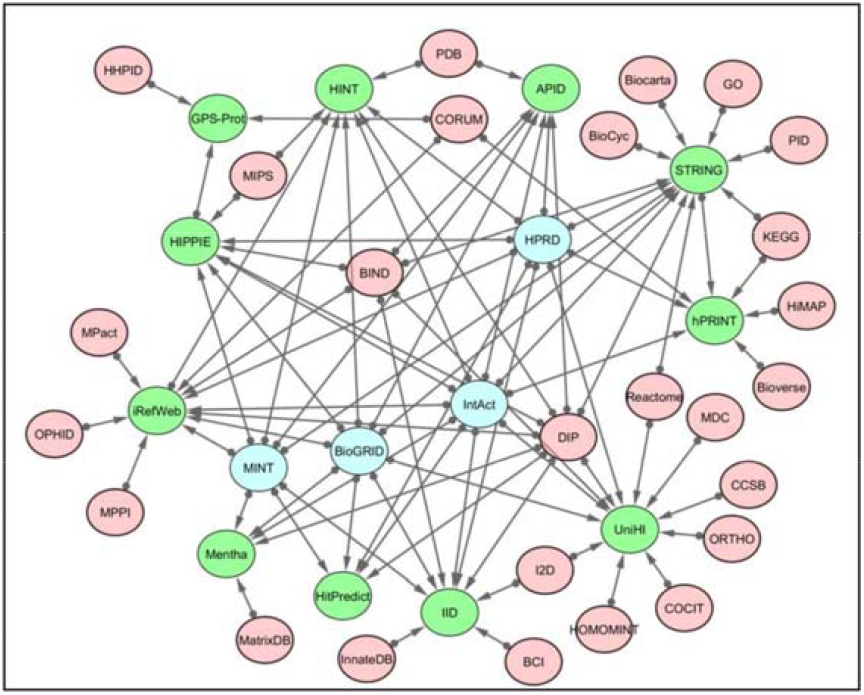
A schematic representation of data flow among primary and secondary protein interaction databases. Green nodes indicate secondary databases and blue nodes indicate primary databases considered in the current study. Data sharing across the databases is shown by edges. The arrows indicate the direction of data flow.

A total of 108 genes including 80 tissue specific (20 genes per tissue), 20 ubiquitous and 28 disease-associated genes (5 genes per disease except for cystic fibrosis, where only 3 genes were found to be associated) were selected for the collection of PPI data. The list of genes is provided in **Supplementary data, Table S2**.

### Comparison of the database-outputs using tissue-specific, ubiquitous and disease associated genes

For all the gene-sets (ubiquitous, tissue-specific and disease associated), hPRINT retrieved highest (61%) ‘total’ interactions. STRING-API and STRING-Web retrieved 46.3% and 20%, respectively, while IID obtained 13.7% PPIs compared to other databases. A combination of STRING-API and hPRINT ensured a yield of ~86% of all possible (total) PPIs, which increased to 90% after including PPIs from IID. For ubiquitous and cardiomyopathy gene-sets, however, iRefWeb showed higher yield (21.5 and 11.8% respectively) than IID (16.7 and 8.6% respectively). On the other hand, for the ‘EV’ interactions, overall, STRING provided the highest yield (STRING-API: 56.9%; STRING-Web: 38.8%) followed by UniHI (31.2%) and iRefWeb (27.9%). However, again, for the ubiquitous gene-set, iRefWeb seemed to provide a little better yield (47% of the ‘EV’ PPIs) than UniHI, which covered 20.5% of ‘EV’ PPIs. Interestingly, for Alzheimer’s associated genes, many databases (e.g., APID, HIPPIE, BioGRID) covered more than 40% of ‘EV’ interactions, whereas, STRING covered only 15.7% of the total possible interactions. Further, HIPPIE showed better yield-coverage for the kidney tissue and lung cancer associated gene-sets (12.8 and 22.9% respectively) compared to iRefWeb. The yield-coverage of hPRINT database for ‘EV’ interactions was very low (6.4%). Primary databases such as, HuRI, MINT, and HPRD, each covered less than 6% of total as well as experimental interactions. Among the 5 primary databases (BioGRID, HPRD, MINT, IntAct, and HuRI) showed BioGRID to have highest yield-coverage for both total and ‘EV’ interactions.

The relative percentage of ‘total’ PPIs for the ubiquitous gene-set was comparatively higher in all the databases, except STRING, hPRINT, HitPredict and HuRI. STRING-Web, hPRINT, and HuRI retrieved slightly more interactions for tissue specific gene-sets, whereas, STRING-API and HitPredict were slightly better for disease specific genes. When experimental PPIs within each database were compared, the relative percentage of PPIs for disease specific genes was higher in all the databases, except in HuRI, IntAct, hPRINT, STRING-API, and iRefWeb. The HuRI, IntAct, hPRINT and STRING-API databases retrieved slightly higher percentage of PPIs for tissue-specific genes, whereas, iRefWeb was relatively better for ubiquitous gene-set. The average interaction yield-coverage for ubiquitous, tissue-specific and disease gene-sets are provided in **Fig 3** and the detailed results are provided in **Supplementary data, Table S3 & S4**.

**Fig 3:**
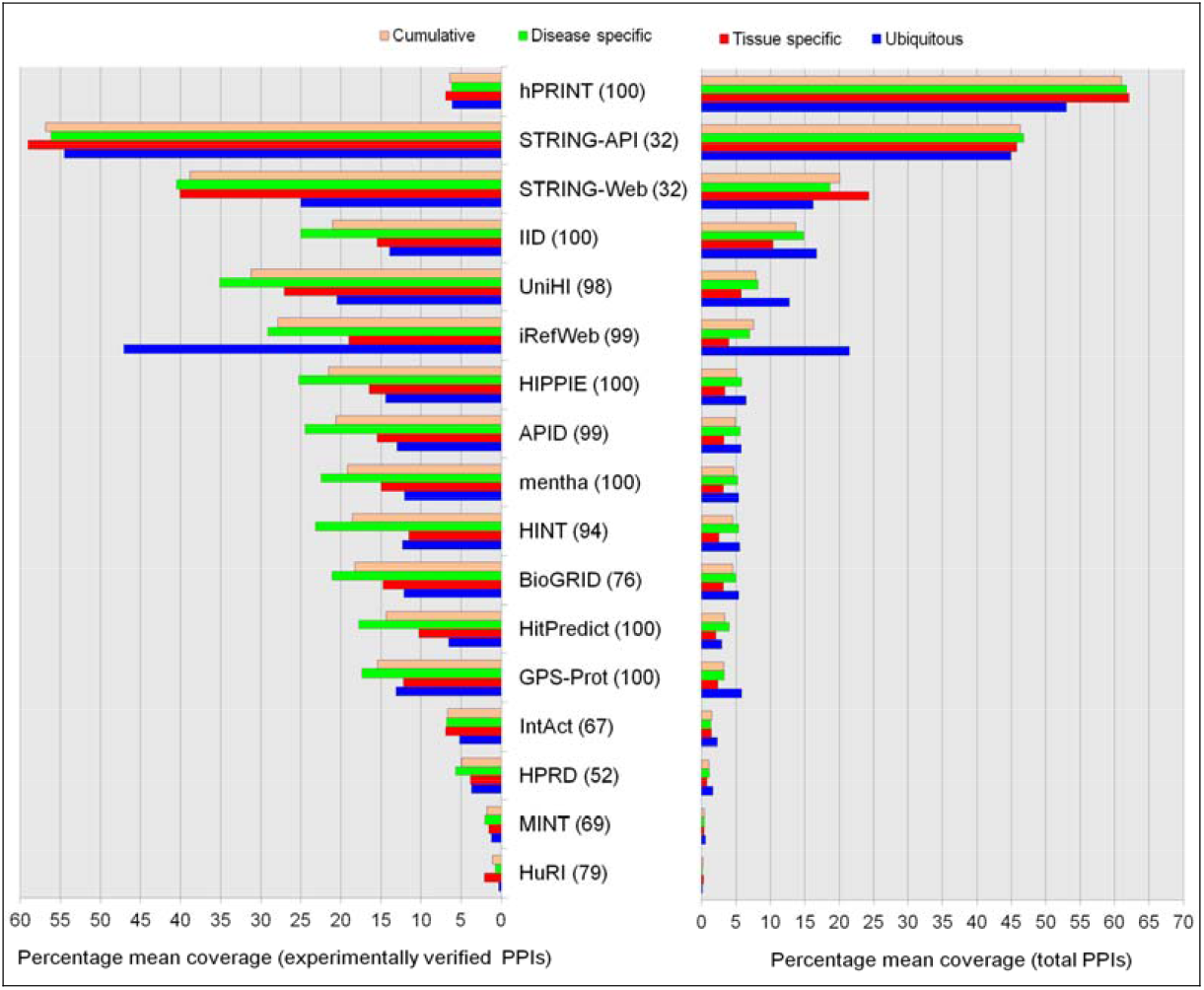
Mean yield-coverage of protein interactions across databases for ubiquitous, tissue specific, and disease associated gene-sets. The databases were manually queried with individual gene/protein names and list of interacting proteins were obtained. A union list of non-redundant interactions for each gene-set across all the databases was considered as reference for calculating the yield-coverage of each database. The yield-coverage for ‘total’ and ‘experimentally verified’ protein interactions was calculated independently by considering the reference data for ‘total’ and ‘experimentally verified’ interactions, respectively. The number in the parenthesis next to the database name indicates its usage frequency rank. More details on the yield-coverage of individual gene-sets can be found in **Supplementary data, Table S3 and S4**.

The comparison using well and less-studied genes overall showed a similar trend. The hPRINT database had highest yield-coverage (60.5 and 55.2% respectively), followed by STRING (STRING-API: 46% and 43.4% respectively; STRING-Web: 19% and 36.1% respectively) and IID (13.4% and 6.3% respectively) for the total interactions, **Supplementary data, Table S5**. When ‘EV’ interactions were compared for well-studied gene-sets, STRING showed highest yield (STRING-API: 54.2%; STRING-Web: 33%) followed by UniHI (31%) and iRefWeb (27.7%) databases. However, for the less studied genes, APID and BioGRID with a mean yield-coverage of 24.3% and 24.1% respectively followed STRING (STRING-API: 53.4%; STRING-Web: 43.9%), **Supplementary data, Table S6**.

Comparison of ‘total’ exclusive interactions across databases for the gene-sets showed hPRINT to have highest mean yield-coverage (39.9%) followed by STRING-API (25.4%) and IID (3.8%), **Fig 4**. For ubiquitous and cardiomyopathy genes, however, iRefWeb (9.5% and 6.7% respectively) and for Alzheimer’s disease, HitPredict (7.2%) provided more number of exclusive interactions than the IID database, **Supplementary data, Table S7**. When ‘EV’ exclusive interactions were compared across the databases, the trend was found to be similar to that of all ‘EV’ interactions, where STRING-API showed highest mean yield-coverage (44.3%) followed by iRefWeb (10.5%), and UniHI (9.8%) databases (**Fig 4**). Again for Alzheimer’s disease, HitPredict showed much better yield-coverage (22.3%) than the other databases. For cystic fibrosis gene-set, IID showed better yield-coverage (4.3%) than iRefWeb, **Supplementary data, Table S8**. Further, the exclusive PPIs from the STRING-Web interface, using selected gene-sets showed a similar result to that of STRING-API. On an average, hPRINT showed highest (44.7%) exclusive hits followed by STRING-Web (9.7%), although, as expected, the percentage of exclusive interactions from STRING-Web was comparatively less than those from STRING-API. When ‘EV’ interactions were considered, STRING-Web showed highest (22.8%) exclusive PPIs followed by iRefWeb (14.1%) and UniHI (8.2%) databases, **Supplementary data, Table S9**. The complete list of protein interactions collected for each gene can be accessed from http://www.startbioinfo.com/PPIs_Individual_Genes.zip.

**Fig 4:**
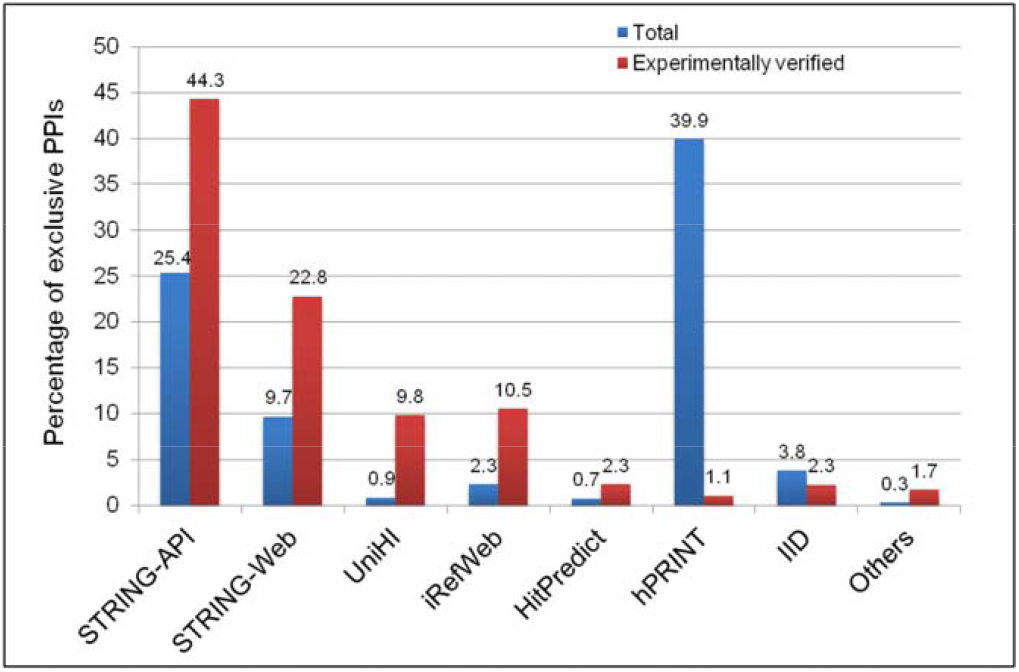
Mean yield-coverage of exclusive protein interactions across databases for the gene-sets. STRING-Web results are based on the protein interactions of selected gene-sets (ubiquitous, kidney, breast cancer, lung cancer, Alzheimer’s disease, and cardiomyopathy). ‘Others’ include, mentha, APID, HIPPIE, GPS-Prot, BioGRID, HPRD, IntAct, MINT, HINT, and HuRI databases. The exclusive interactions represented for UniHI, iRefWeb, HitPredict, hPRINT, IID, and Others are based on the comparison with STRING-API resulted PPIs and not STRING-Web. A union list of non-redundant interactions across all the databases was considered as reference for calculating the exclusive yield-coverage of each database. More details on the exclusive interactions for individual gene-sets can be found in **Supplementary data, Table S7, S8, and S9**.

### Comparison of complete protein interaction data across databases

Complete human protein interaction data comparison showed hPRINT to have highest number of total interactions (58.1%) followed by STRING (44.7%) and IID (10%). When the number of proteins corresponding to total interactions was compared, STRING database was found to have the highest coverage. hPRINT, followed by STRING and IID had highest number of mean ‘total’ interactions per protein. In case of ‘EV’ interactions, STRING covered 79.3% of the all possible interactions, followed by APID (16.9%) and HIPPIE (16.5%). A similar trend was observed for the average number of ‘EV’ interactions per protein. APID provided highest number of proteins corresponding to the ‘EV’ interactions, followed by HitPredict and HIPPIE databases. Among the five primary databases considered, BioGRID showed highest content-coverage for both interactions as well as proteins. APID database was found to have highest number of PPIs, followed by HIPPIE and HitPredict when the databases containing only experimentally verified interactions were considered. Among the databases storing both ‘EV’ and predicted protein interactions, iRefWeb had highest percentage of ‘EV’ interactions followed by UniHI, whereas, hPRINT had the least. The details on the content-coverage of interactions and proteins for different databases are provided in **Table 2** and **Fig 5**.

**Table 2:** Summary of complete protein interaction data downloaded from the databases.

| Database name | Number of total PPIs (EV PPIs) | Percentage EV PPIs | Total PPIs |  | EV PPIs |  |
| --- | --- | --- | --- | --- | --- | --- |
|  |  |  | Number of proteins | Avg. number of PPIs/protein | Number of proteins | Avg. number of PPIs/protein |
| STRING† | 4,056,217 (1,620,507) | 40.0 | 18,523 | 219 | 16,366 | 99 |
| UniHI† | 359,772 (267,393) | 74.3 | 17,311 | 21 | 16,635 | 16 |
| iRefWeb† | 168,549 (145,874) | 86.5 | 15,266 | 11 | 14,989 | 10 |
| Mentha | 270,130 | -- | 16,477 | 16 | -- | -- |
| APID | 345,965 | -- | 17,473 | 20 | -- | -- |
| HIPPIE | 337,892 | -- | 17,283 | 20 | -- | -- |
| BioGRID* | 260,308 | -- | 16,682 | 16 | -- | -- |
| HPRD* | 39,001 | -- | 9,608 | 4 | -- | -- |
| HitPredict | 262,317 | -- | 17,371 | 15 | -- | -- |
| IntAct* | 143,973 | -- | 14,689 | 10 | -- | -- |
| MINT* | 17,265 | -- | 6,215 | 3 | -- | -- |
| HINT | 121,957 | -- | 14,433 | 8 | -- | -- |
| hPRINT† | 5,272,669 (93,149) | 1.8 | 18,192 | 290 | 11,167 | 8 |
| HuRI* | 25,188 | -- | 7,944 | 3 | -- | -- |
| IID† | 905,499 (279,529) | 30.9 | 17,916 | 51 | 16,447 | 17 |

†Databases storing both experimentally verified and predicted protein interactions; \*Primary protein interaction databases. Note: Non-redundant PPIs and proteins of each database are represented. EV: Experimentally Verified

**Fig 5:**
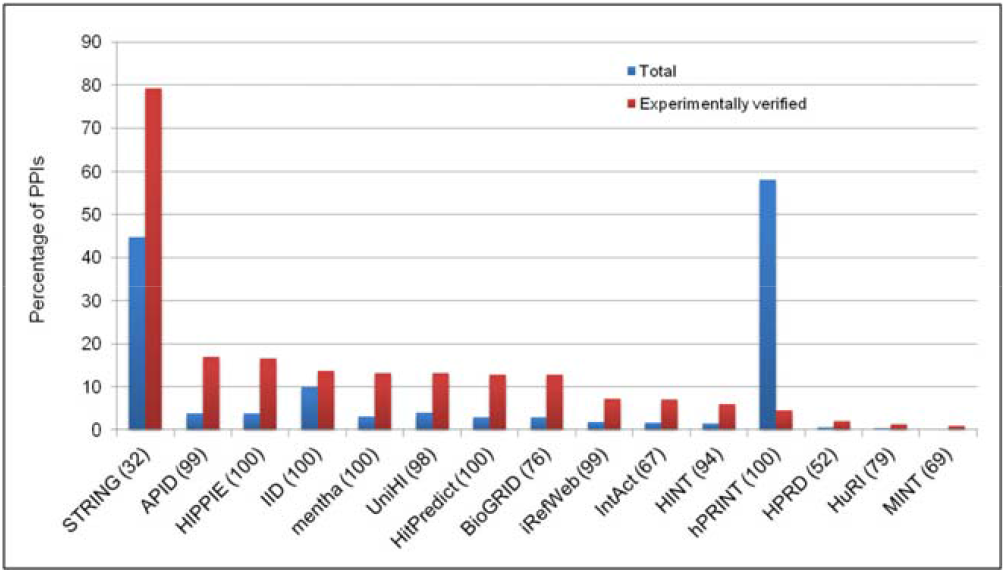
Coverage of protein interactions across the databases based on complete downloaded data. A union list of non-redundant interactions across all the databases was considered as reference for calculating the content-coverage of each database. The content-coverage for ‘total’ and ‘experimentally verified’ interactions was calculated independently by considering reference data for ‘total’ and ‘experimentally verified’ interactions respectively. The number in the parenthesis next to the database name indicates its usage frequency rank.

When the databases were compared for their exclusive contribution of proteins, hPRINT showed highest coverage followed by UniHI (**Fig 6**). The hPRINT database was also found to have highest content-coverage for ‘total’ exclusive interactions (46.5%) followed by STRING (33.1%) and IID (3.1%). However, STRING provided most number of ‘EV’ exclusive interactions (70.6%) and was followed by UniHI (5%). Six databases, hPRINT, STRING, UniHI, IID, APID, and HIPPIE together covered 99.7% of ‘total’ and 98.9% of ‘EV’ exclusive interactions (**Fig 7**).

**Fig 6:**
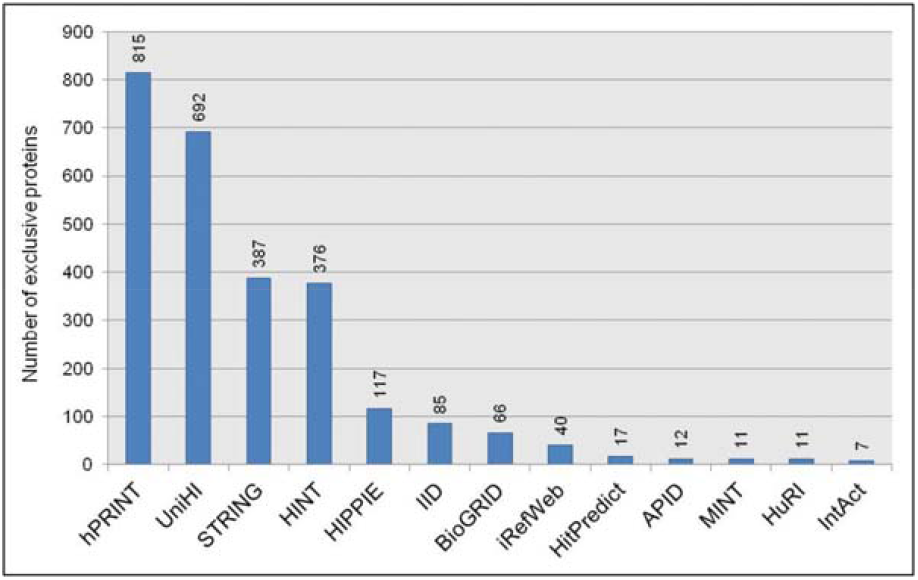
Number of exclusive proteins contributed by each database. A non-redundant list of proteins was obtained from the interaction pairs downloaded from each database and compared across all the databases to obtain exclusive number of proteins. Only databases with exclusive contribution are shown in the figure (mentha and HPRD had no exclusive contribution). Numbers corresponding to each bar represent the number of exclusive proteins in the database.

**Fig 7:**
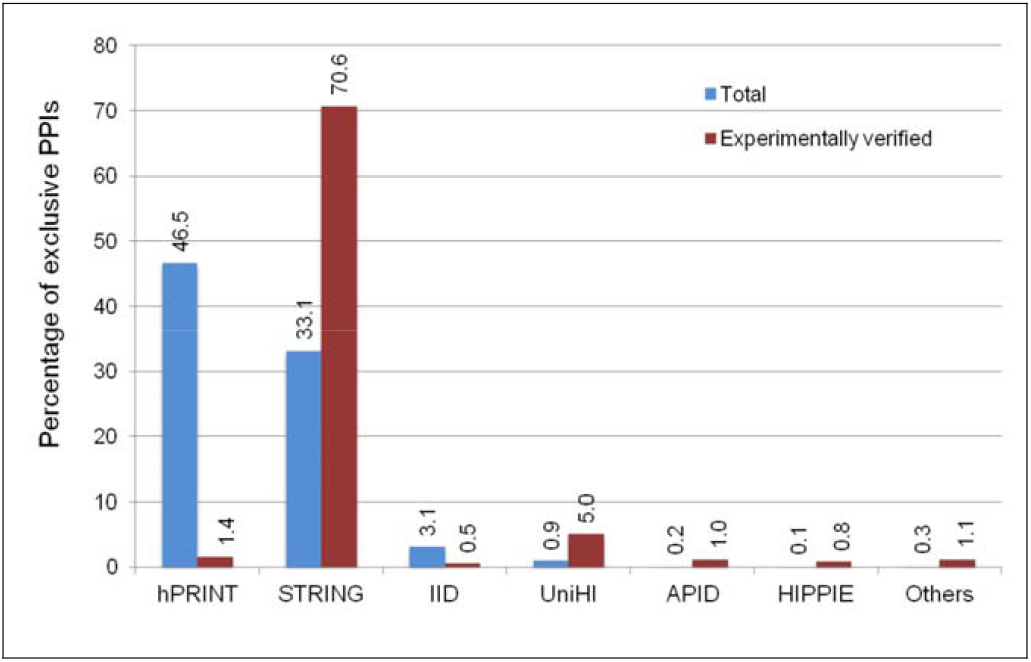
Coverage of exclusive protein interactions in the databases, based on the analysis of complete downloaded data. The interaction pairs extracted from the downloaded data were compared across the databases to obtain exclusive interactions in each database. The comparison was performed with both the possible combinations of interaction pairs (P1 – P2 and P2 – P1). A union list of non-redundant interactions across all the databases was considered as reference for calculating the content-coverage of each database for ‘total’ and ‘experimentally verified’ interactions independently. ‘Others’ include iRefWeb, mentha, BioGRID, HPRD, HitPredict, IntAct, MINT, HINT, and HuRI.

The HPRD database did not contribute any exclusive interaction. IntAct had the least number of exclusive interactions followed by MINT for both ‘total’ and ‘EV’ PPIs. Among the primary databases, BioGRID provided highest number of exclusive interactions (49.5%) followed by IntAct (17.2%) corresponding to 69.5% and 24.2% of all the exclusive interactions, respectively. MINT contributed least number of exclusive interactions among the primary databases (**Fig 8**).

**Fig 8:**
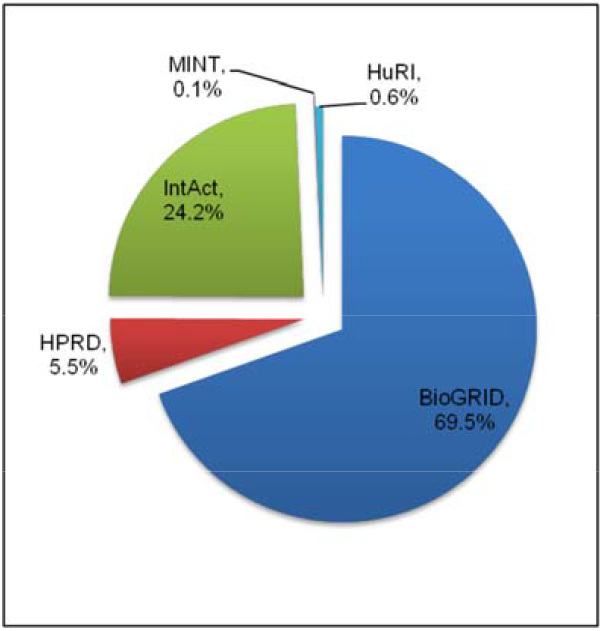
Coverage of exclusive protein interactions among the primary databases, based on the analysis of complete downloaded data. The interaction pairs extracted from the downloaded data were compared across the primary databases to obtain exclusive interactions. The comparison was performed with both the possible combination of interaction pairs (P1 – P2 and P2 – P1). A union list of non-redundant interactions across all the primary databases was considered as reference for calculating the exclusive interactions contributed by each database.

The comparison of databases based on number of ‘EV’ PPIs per protein showed that ~90% of proteins in both MINT and HuRI databases had 10 or fewer interactions. At the same time, 34% of the proteins (5,580 proteins) in STRING database had at least 100 interacting partners (**Fig 9**). It was interesting to note that certain proteins have high number of interactions, with ubiquitin C (UBC), ubiquitin B (UBB) and ubiquitin D (UBD) on top of the list having 11,849, 4,435, and 3,928 PPIs, respectively (**Fig 10**).

**Fig 9:**
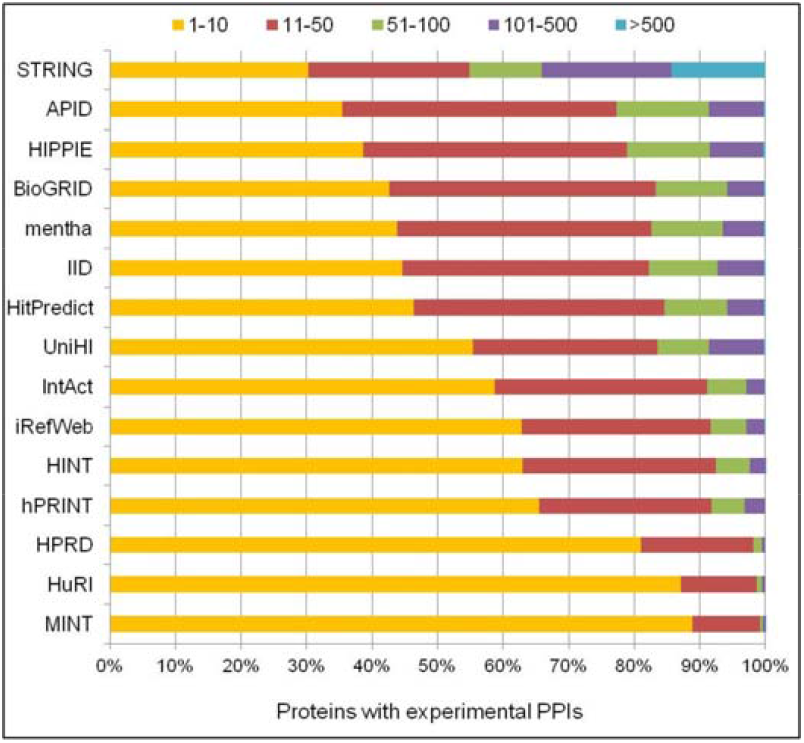
Number of experimentally verified protein interactions per protein. A non-redundant list of interaction pairs was obtained for each database (P1-P2 and P2-P1 were treated same and only one pair was retained). Number of interactions per protein in each database was derived by counting the number of times a protein is present in the non-redundant list of each database.

**Fig 10:**
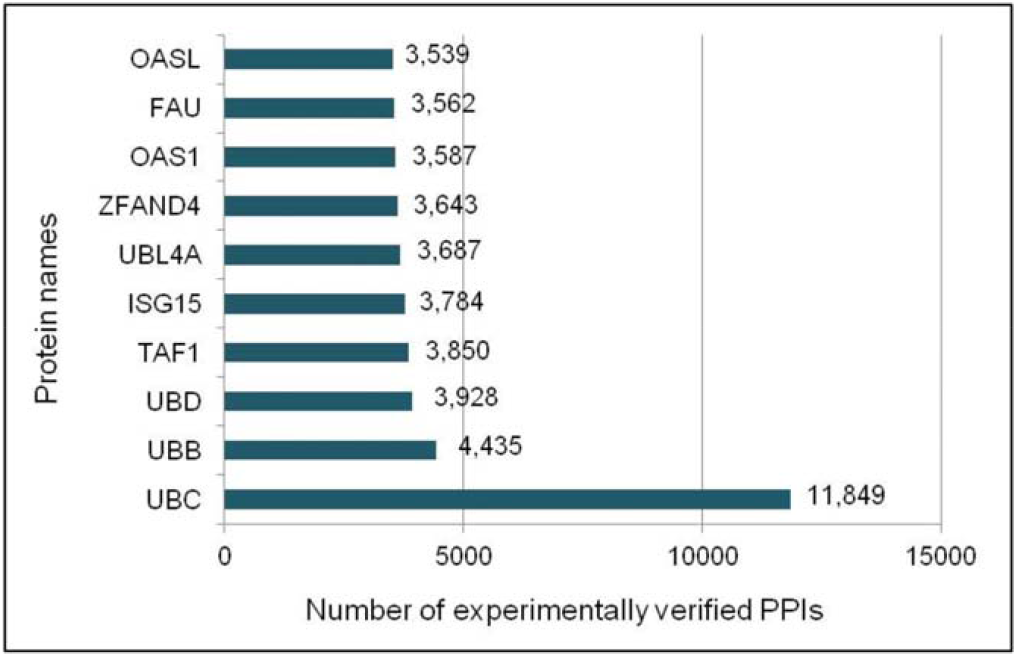
Top 10 proteins based on number of experimentally verified protein interactions. A non-redundant list of experimentally verified interaction pairs was obtained by combining PPIs downloaded from all the databases (P1-P2 and P2-P1 were treated same and only one pair was retained). Number of interactions per protein was derived by counting the number of times a protein is present in the combined list. Numbers corresponding to each bar represent the number of PPIs for the protein.

## Discussion

The first time compilation of 375 PPI resources, followed by short-listing and basic comparison of 125 relevant resources for human PPIs and then a systematic comparison of 16 selected databases has provided many new insights. The study shows specific advantages of different databases for specific purposes and indicates benefits of combinatorial usage of certain databases in some cases, for the first time. These observations and resulting suggestions are different from some of the earlier studies. A study [8] on comparative analysis of six protein interaction databases showed IntAct to be the most comprehensive database in terms of number of PPIs as well as species. The authors observed that the pairwise overlap among the considered databases was only up to 75%. Further, to increase the number and quality of protein interaction data, the study suggested a submission requirement of the PPI data prior to publication. In the current study, IntAct contributed <10% of all the possible interactions, and a very less percentage (<1 %) of exclusive interactions. Another study [9] showed that HPRD [26] covered highest number of proteins, interactions, and literature citations. However, it was found to be low in terms of contents as well as output in our study. The lower coverage of IntAct and HPRD could be because of the consideration of meta-databases in the current study, unlike the previous studies and/or lack of updates (e.g., HPRD was last updated in 2010). However, despite their low coverage, these two PPI databases seem to be very popular as indicated by their usage frequency, which were next to STRING (**Fig 3**). Lack of the meta-databases until recently combined with preference for manually curated data and observations from an earlier comparative study [9] may have prompted researchers to use these databases. Among the primary databases, HPRD retained the top position in terms of usage frequency, even though BioGRID shows highest coverage (69.5%) of PPIs according to our study (**Fig 8**). Such observations stress the need for detailed comparative analysis as the current one.

Overall, hPRINT and STRING showed highest coverage for total and ‘EV’ interactions respectively, for both gene-based queries and complete PPI data. These databases also contributed maximum number of exclusive interactions, which makes them a suitable choice for any research involving PPIs. Moreover, these are meta-databases that cover data from multiple resources, and also predict interactions using several computational methods. Though, hPRINT provided highest number of total interactions, the coverage for the ‘EV’ PPIs was found to be <10%. This is perhaps due to the variation in the annotation of ‘EV’ PPIs in hPRINT compared to the rest. Even though hPRINT integrates data from multiple resources including STRING (which had highest coverage for the ‘EV’ interactions), the interactions referred to be experimentally verified in STRING are not categorized in case of hPRINT. The STRING database which showed a very good coverage of PPIs in the current study has also been found to be frequently used (based on the UFR) by the scientific community, probably due to its publication in well-known journal(s) and data update frequency. Whereas, hPRINT being in the top position for its coverage found to be at the bottom in terms of its usage (**Fig 3**), probably because it is comparatively new. Further, the UFR specifically indicated that the usage of some of the databases has been limited even though they have high coverage. One of the reasons behind this could be the lack of awareness among the scientific community about the availability of such resources with better coverage. Hence, compilation and periodical comparative studies of the PPI databases and tools are a need of the hour, which can equip biologists with latest developments and help them in choosing the right resource.

hPRINT provided a good coverage of total interactions, but it does not support any visualization of interactions, or integrate additional data for the proteins, such as pathways, domains. The output of hPRINT is just the interactions in a tabular format, with their evidence scores. On the other hand, STRING, along with a good coverage of interactions, offers visualization of the interaction network. It also supports functional enrichment of the network proteins using pathways, domains, and Gene Ontology annotations. The data in STRING is updated on a regular basis, and the complete data can be downloaded in various formats aiding for an easier integration into any new tool. It should be noted that STRING has limited the number of interactions to be displayed in the web-browser to 500. However, as suggested by the database, the API can be used to obtain all the interactions for a particular protein. Further, the lower yield of ‘EV’ protein interactions in STRING database for Alzheimer’s disease can be attributed to APP and A2M genes, which retrieved 3-5 times lesser number of PPIs than most other databases. In the current study, both STRING-Web and STRING-API were independently queried with the genes to obtain the interactions. Although STRING-Web provided less number of interactions compared to STRING-API, in terms of protein interaction yield, the former was found to be still better than most of the other considered databases for both ‘total’ and ‘EV’ interactions as well as for exclusive contribution. Hence, a user can obtain good number of interactions, if not all, by using the web-based interface of STRING. The decision of using STRING-API versus STRING-Web can be made based on the research goal. When all the interactions for a particular protein are required, it may be better to use the API; whereas the web-based interface would be very useful to obtain and visualize a network of high confidence interactions only. The results show that the protein interaction coverage of primary databases was comparatively low both in terms of ‘all’ and ‘exclusive’ contribution. This is expected due to two reasons: a) the primary databases collect interaction through literature curation, which is a manual process and takes enormous amount of time to populate the database; b) most of the secondary databases integrate PPIs from multiple primary databases, and hence have a higher coverage. No exclusive contribution (based on complete data comparison) by HPRD database could be because the PPI data has not been updated since April 2010.

The IID database, despite having a lower coverage, can be particularly useful in obtaining PPIs along with the expression status of interacting proteins in different tissues. Similarly, HIPPIE database provides an option to filter the interacting proteins based on their tissue specificity. TissueNet [35] is another database (not considered in the current study) similar to IID, which provides tissue associated protein interactions. The IID database collects the gene expression data from microarray studies, whereas, the expression data in HIPPIE and TissueNet has been obtained from GTEx, a RNA sequencing based expression database. The protein expression data in both IID and TissueNet has been integrated from Human Protein Atlas. Currently, except Wiki-Pi [36] and IntAct, no other protein interaction database provides disease associated PPIs. However, the association in Wiki-Pi is based on Gene Ontology (GO), functional, or pathway annotations of the interacting proteins, which have been integrated from HPRD and BioGRID databases. On the other hand, IntAct provides a precompiled list of interactions of proteins associated with diseases, such as cancer, diabetes, Alzheimer’s, Parkinson’s. ComPPI [37] is another resource that integrates GO annotations related to sub-cellular locations of interacting proteins, greatly enhancing the reliability of the interactions. Interactome3D [38] is a web-server useful for structural annotation of the protein interaction networks. When queried with an interaction, it can find the available structural data for both the interacting partners as well as for the interaction itself.

While performing the comparative analysis, we also noted the shortcomings of different databases from a user’s perspective, which we hope will be resolved in due course of time or in the future versions of the databases, making them more user-friendly and reliable. For example, GPS-Prot provided no result for some proteins (e.g., ESR1) even after taking hours to load the interaction network, making it difficult for us to judge whether the database has any interaction for the specific protein. In hPRINT database, the download option for the output retrieved for gene-based queries resulted in a terminated file with incomplete list of interacting partners, and hence, interactions were collect manually by copying them from the result page. The STRING database provides maximum of 500 interactions through the web based query, probably because of the graphical interaction network. Hence, the user has to use API or download the complete interaction data to obtain all the available interactions for a protein of interest. Further, this limitation can be clearly indicated in the web-page and can be overcome by providing an option to download all the available interactions for the query protein, while limiting the graphical network to maximum of 500 interactions. IID database when queried with gene/protein name in small letters (e.g., tp53 instead of TP53) does not provide any interaction. This limitation was evaluated using multiple gene/protein names and different browsers.

The whole study has been performed in a span of a few years, where the compilation of protein interaction resources took most of the time. The collection of interactions and comparative analysis of the databases, however, was performed within a year. Therefore, variations may be expected in the number of interactions for some databases based on their update status. For example, mentha updates the interactions every week, whereas, HINT claims to update its interaction data every night, making it practically impossible to get the exact coverage of these databases at a given time point. However, based on our preliminary studies, we believe that this will not affect the results drastically and the overall trend in the coverage of the databases will remain same at least in the near future. Further, for the comparison of complete interaction data, we have considered only binary interactions, while complex interactions were excluded. Also, we did not assess the accuracy of the interactions provided by different databases, as it is beyond the scope of our study. However, study by Turinsky et al., [7] quantified the agreement between curated interactions shared across nine major public databases. Their results indicated that on an average, two databases fully agree on approximately 40% of the interactions and 60% of the proteins curated from the same publication. They also showed that the variation can be mainly because of the difference in assignments of organism or splice isoforms, different organism focus, and alternative representations of multi-protein complexes, among others.

## Conclusions

Realizing the need for periodic comparative analysis of protein-interaction databases, we compiled 375 resources that are directly or indirectly useful in studying protein-protein interactions. Apart from basic comparison of 125 resources that are likely to be more relevant for researchers, this study completes one of the most elaborate evaluations of 16 carefully selected databases from a user’s perspective. The results showed that hPRINT, followed by STRING, would be ideal for obtaining maximum of the protein interactions. Together, they contain ~91% of the ‘total’ interactions as indicated from the analysis of complete downloaded data. When the focus is on only Experimentally Validated (EV) interactions, STRING could be the choice (~79% ‘EV’ PPIs), though a combinatorial use of STRING and UniHI would ensure a slightly higher content-coverage of ‘EV’ interactions (~84%). STRING-Web, although provides less number of interactions than its API counterpart, is still better than most of the other databases in terms of coverage of all and exclusive interactions. Our results also conclude that a universal rule cannot be applied for selecting the databases for gene-based searches of PPIs. Among the five primary PPI databases considered, BioGRID provided highest coverage. The current compilation of resources, review of several features of the important databases, and the findings of the comparative study can help researchers in making an objective-oriented selection of suitable protein-protein interaction database(s), particularly for human species. The current study may also pave new components in the methods for comparing bioinformatics resources especially from a user’s perspective.

## Acknowledgements

The authors thank colleagues at BdataA and Shodhaka Life Sciences Pvt. Ltd. for assistance in gathering some of the PPI data and analysis.

## Supplementary Information

**S1 Text.** Protein interaction databases selected for detailed comparative analysis

**S2 Text.** Selection of ubiquitous, and testis, kidney and uterus specific genes

**S1 Table.** Conversion of gene/protein identifiers in complete downloaded data files to official gene symbols

**S2 Table.** Gene-sets selected for the comparison of protein interaction databases

**S3 Table.** Coverage of ‘total’ protein interactions for all gene-sets

**S4 Table.** Coverage of ‘experimentally verified’ protein interactions for all gene-sets

**S5 Table.** Coverage of ‘total’ protein interactions for well and less studied gene-sets

**S6 Table.** Coverage of ‘experimentally verified’ protein interactions for well and less studied genes

**S7 Table.** Coverage of ‘total’ exclusive protein interactions for all gene-sets

**S8 Table.** Coverage of ‘experimentally verified’ exclusive protein interactions for all gene-sets

**S9 Table.** Coverage of exclusive protein interactions using STRING-Web for selected gene-sets

Author contributions
Rationale, Study-design and Guidance: **KKA**
Data compilation, Standardization, Comparisons, and Coordination across the team: **AKB** and **SD**.
Data compilation and Comparisons: **KT, SN**, and **SO**.
Data validation: **AKB, SD**, and **SO**
Gene list preparations: **KB, KT, AKB**
Data downloading and scripts to compare: **AKB** and **DD**
General supervision and comments: **KTH**

