## Supplementary Information for "How helpful are the protein-protein interaction databases and which ones?"

**Supporting Information**

**S1 Text. Protein interaction databases selected for detailed comparative analysis**

A total of 16 protein interaction databases were selected from an initial list of 375 PPI resources (**Startbioinfo:** <http://startbioinfo.com/cgi-bin/simpleresources.pl?tn=PPI_AR>**)** for the detailed comparative analysis of human protein interactions. A brief description of the selected databases along with their features is provided below.

**STRING** (*Search Tool for the Retrieval of INteractinG proteins:* [*http://string-db.org*](http://string-db.org/))**:** A meta-database that stores both known and predicted interactions. The known interactions are collected from various protein interaction and pathway databases (BIND, DIP, BioGRID, HPRD, IntAct, MINT, PID, Biocarta, BioCyc, GO, KEGG, and Reactome). The interactions can be visualized in a network with different modes. The evidence mode shows multiple colored edges between the nodes indicating the type of evidence, whereas, in confidence mode the thickness of the edges changes based on the confidence score. In case of molecular action mode, the edges indicate the function of the interactions. The database uses methods, such as gene neighborhood, co-expression, gene fusion, and co-occurence to predict the interactions among proteins. It also uses text-mining to collect the interactions from literature. Each interaction is assigned a score based on the type and number of evidences present. The score ranges from 0 to 1, where 0 indicates the lowest and 1 indicates the highest threshold. The database also results in a network among the list of the queried proteins when ‘multiple proteins’ option is used. The STRING database not only provides interactions but also aids in the functional enrichment analysis of the network proteins.

**UniHI** (*Unified Human Interactome:* [*http://www.unihi.org*](http://www.unihi.org/))**:** A meta-database that stores both known and predicted interactions from various databases (MDC, CCSB, HPRD, BioGRID, BIND, DIP, IntAct, Reactome, HOMOMINT, COCIT, ORTHO and I2D). It accepts gene or protein identifiers from different organisms as query and results in physical and regulatory interactions in the human interactome. The database supports visualization of the obtained results and offers multiple filtering options, such as based on source of interactions, evidences. The drug targets can also be easily analyzed in the resulting interaction network. Expression data by the user can be uploaded and tissue-, process- or disease-specific networks can be created using the integrated tools. The database also provides tools for exploring the functional relevance of molecular networks.

**Mentha** ([*http://mentha.uniroma2.it/*](http://mentha.uniroma2.it/)): A meta-database which collects interaction data from manually curated protein-protein interaction databases (MINT, IntAct, DIP, MatrixDB, and BioGRID) that have adhered to the International Molecular Exchange (IMEx) consortium. The database also offers a series of tools to analyze selected proteins in the context of a network of interactions. The interaction data is updated every week to generate a consistent interactome. It also assigns each interaction a score based on the supporting evidences.

**hPRINT** ([*http://print-db.org/hprint-web/*](http://print-db.org/hprint-web/)): It uses a combination of random forest and Bayesian learning approaches in order to integrate various evidences (text mining, genetic relationships, evolutionary information, and domain profiles) for predicting protein interactions and integrating the predictions with known information. The interaction data is collected from different sources, including known and predicted interaction databases (STRING, HiMAP, Bioverse, KEGG, HPRD, IntAct, and CORUM) and published articles.

**APID** (*Agile Protein Interactomes DataServer:* [*http://apid.dep.usal.es/*](http://apid.dep.usal.es/))**:** A meta-database that provides a comprehensive collection of experimentally verified protein interactions from BIND, BioGRID, DIP, HPRD, IntAct, MINT, and PDB. It includes a visualization web-tool that allows the construction of sub-interactomes using query lists of proteins of interest and the visual exploration of the networks. The visualization tool also helps in an interactive selection of the interactions based on various network properties such as, reliability of the edges and functional annotations of the nodes in the network.

**HIPPIE** (*Human Integrated Protein-Protein Interaction rEference:* [*http://cbdm-01.zdv.uni-mainz.de/~mschaefer/hippie/*](http://cbdm-01.zdv.uni-mainz.de/~mschaefer/hippie/))**:** A meta-database that collects experimentally verified protein interactions from IntAct, BioGRID, HPRD, DIP, MINT, BIND, MIPS, and literature. A core component of the database is the confidence scoring of interactions based on the supporting evidences. The score is calculated as a weighted sum of the number of studies in which an interaction was detected, the number and quality of experimental techniques used and the number of non-human organisms in which an interaction was reproduced. The network construction mode can be used to graphically visualize the interaction network among the query proteins. The network can also be filtered based on interaction type, tissue specificity, or functional annotations.

**GPS-Prot** ([*http://www.gpsprot.org*](http://www.gpsprot.org/))**:** A web-based platform that integrates different HIV interaction data types as well as interactions between human proteins derived from publicly-available experimentally verified databases (HIPPIE, CORUM, HHPID), and literature. The web-based visualization tool allows users to create comprehensive and integrated HIV-host networks. The tool can group proteins into functional modules or protein complexes, and helps in generating more intuitive network representations. The users can also upload their own data and perform integrated analysis.

**BioGRID** (*Biological General Repository for Interaction Datasets:* [*http://thebiogrid.org*](http://thebiogrid.org/))**:** A public database that stores genetic and protein interaction data from model organisms as well as humans. The interactions are curated from high-throughput datasets as well as from individual focused studies. Complete coverage of the entire literature is maintained for budding yeast (*Saccharomyces cerevisiae*), fission yeast (*Saccharomyces pombe*) and thale cress (*Arabidopsis thaliana*).

**HPRD** (*Human Protein Reference Database:* [*http://www.hprd.org*](http://www.hprd.org/))**:** A database focused on human protein related data such as, posttranslational modifications, disease associations, tissue expression, sub-cellular localization, and protein interaction. Most of the protein related data, including protein interaction has been obtained from primary literature by manual curation.

**HitPredict** ([*http://hintdb.hgc.jp/htp/*](http://hintdb.hgc.jp/htp/))**:** A meta-database of experimentally verified protein interactions collected from 5 primary interaction databases (IntAct, BioGRID, HPRD, DIP, and MINT). The interactions in the database are assigned a reliability score based on the experimental information available and other details such as, structurally known interacting pfam domains, Gene Ontology annotations for the interacting proteins, and homologous interactions.

**IntAct** ([*http://www.ebi.ac.uk/intact*](http://www.ebi.ac.uk/intact))**:** A database of experimentally verified protein interactions derived from literature curation or direct user submissions. The interactions can be restricted to specific field, such as organism, experimental method, and interaction type. Further, the database provides interaction between protein and chemical molecules collected from ChEBI chemical database. IntAct also stores curated protein interaction datasets for a few diseases, such as cancer, Diabetes, Parkinson’s disease, which can be downloaded in the PSI-MI compliant format.

**MINT** (*Molecular INTeraction:* [*http://mint.bio.uniroma2.it*](http://mint.bio.uniroma2.it/)): A public repository of experimentally verified protein interactions collected from literature by expert curators. MINT is one of the members of the IMEx that allows sharing of curation efforts. Each interaction is assigned a reliability score based on number of studies and types of experiments used. It also provides graphical visualization of the interacting partners. In addition to binary and complex protein interactions, the database also stores other types of functional interactions, including enzymatic modifications of the interacting partners.

**HINT (***High-quality INTeractomes:* [*http://hint.yulab.org/*](http://hint.yulab.org/)): A meta-database of high-quality curated protein interactions collected from 8 interactome resources (BioGRID, MINT, iRefWeb, DIP, IntAct, HPRD, MIPS, and PDB). The interactions are filtered both systematically and manually to remove erroneous and low-quality data. The database contains only manually validated high-throughput experiments and interactions from small-scale studies that have been reported by at least two independent publications in the literature. The database is updated every night.

**IID** (*Integrated Interaction Database:* [*http://ophid.utoronto.ca/iid*](http://ophid.utoronto.ca/iid)): A meta-database of known and predicted eukaryotic protein-protein interactions in 30 tissues of model organisms and humans. It collects the known interactions from 9 major databases (BioGRID, IntAct, I2D, MINT, InnateDB, DIP, HPRD, BIND, and BCI). It also integrates the orthologous protein interactions, and high-confidence computationally predicted interactions from recent studies. The database provides tissue specific protein interactions based on the expression of the interacting partners in specific tissues. It can take input proteins from one organism and can provide tissue-specific interactions in multiple specified organisms by ortholog mapping of the input proteins.

**HuRI** (*Human Reference protein Interactome:* [*http://interactome.baderlab.org/*](http://interactome.baderlab.org/)): A database that stores binary protein interactions by systematically interrogating all pair-wise combinations of predicted human protein-coding genes using yeast two-hybrid (Y2H). The interactions are further validated using multiple orthogonal assays. The database also collects curated set of interactions reported in the literature from small scale studies having at least two different experimental evidences in the original publication.

**iRefWeb** ([*http://wodaklab.org/iRefWeb/*](http://wodaklab.org/iRefWeb/))**:** A meta-database that integrates experimental and predicted protein interaction data from different databases (BIND, BioGRID, CORUM, DIP, IntAct, HPRD, MINT, MPact, MPPI, and OPHID) for multiple species. The interaction data can be filtered based on various criteria, such as the number of supporting publications, the scale of the corresponding studies (high- or low-throughput) or the detection methods used. Further, the interactions can be compared across different databases, to uncover possible inconsistencies, and follow up on them through links to source databases and various supporting evidences.

S1 Table. Conversion of gene/protein identifiers in complete downloaded data files to official gene symbols

| **Database name** | **Gene/protein identifiers in the downloaded file** | **Successful conversion (%) to official gene symbols** |
| --- | --- | --- |
| **HIPPIE** | Uniprot accession | 100 |
| **HitPredict** | Uniprot accession, Uniprot ID, Entrez gene, Ensembl transcript | 99.6 |
| **hPRINT** | Ensembl gene | 97.5 |
| **HuRI** | Ensembl gene, Uniprot ID | 99.5 |
| **IntAct** | Uniprot ID | 97.0 |
| **iRefWeb** | Uniprot ID | 93.1 |
| **MINT** | Uniprot ID | 99.3 |
| **STRING** | Ensembl protein | 95.1 |
| **Average** | | **97.6** |

**S2 Text. Selection of ubiquitous, and testis, kidney and uterus specific genes**

To cover a wide range of case studies, gene-sets related to ubiquitous, and specific to testis, uterus, and kidney tissues were considered for the comparison of protein-protein interaction databases.

The ubiquitous genes were selected from the following study [Chang et al., *PLoS One 2011;6:e22859*]. These genes were further filtered based on their presence in testis [Acharya et al., *BMC Genomics 2010;11:467*], uterus [Bajpai et al., *PLoS One 2012;7:e36776*], and kidney (<http://resource.ibab.ac.in/MGEx-Kdb/>) databases. The corresponding reliability scores for each gene were obtained from these databases. To make them comparable across the databases, the reliability scores in each list were scaled from a range of 0 to 10, where 0 being the lowest and 10 being the highest. The scaled value for each gene was then added across the databases to obtain a cumulative score, and the genes were sorted based on the descending order of scores. Top 100 genes were selected based on the cumulative score and number of citations corresponding to each gene was retrieved from the literature. Top 10 genes and bottom 10 genes based on number of citations were selected as the well studied and less studied genes respectively for the comparative analysis.

For selecting testis specific gene-set, genes transcribed in testis and dormant in kidney and uterus tissues were obtained from the in-house databases. Further, tissue specific genes from two well known databases, TiGER [Liu et al., *BMC Bioinformatics 2008;9:271*] and PaGenBase [Pan et al., *PLoS One 2013;8:e80747*] were obtained. Genes common to all the 5 lists were selected. The reliability scores from testis, kidney, and uterus databases for the common genes were obtained and scaled from 0 to 10 to make them comparable across the three databases. The scaled scores for each gene were then added to obtain the cumulative score, and the genes were then sorted on descending order of scores. Top 100 genes from the list were selected based on the cumulative score and number of citations each gene was retrieved from the literature/Entrez gene database. A total of 10 genes having highest (well studied) and 10 genes having lowest number of citations (less studied) were selected for the comparison of PPI databases.

A similar strategy was employed for the selection of kidney specific genes. The genes transcribed in kidney tissue were screened for their presence in both TiGER and PaGenBase databases, and 90 genes common to all the three databases were obtained. Dormant genes in testis and uterus tissues were not considered for selecting this list, as it resulted in less number of genes than the desired number. Ten well and 10 less-studied genes were then selected based on the strategy used for selecting the testis specific genes.

For selecting uterus specific gene-set, genes common to uterus transcribed list and dormant in kidney and testis associated lists were obtained. Then the reliability scores from each database were scaled from 0 to 10. Further, a similar strategy used for selecting the testis specific genes was used to select 10 well- and 10 less-studied genes for the comparative analysis.

S2 Table. Gene-sets selected for the comparison of protein interaction databases

| **Ubiq** | **Kid** | **Tes** | **Uter** | **LC** | **BC** | **Alzh** | **Diab** | **CM** | **CF** |
| --- | --- | --- | --- | --- | --- | --- | --- | --- | --- |
| SOD1 | SPP1 | PRL | ESR1 | FASLG | RAD54L | HFE | GPD2 | CAV3 | FCGR2A |
| RHOA | EGF | HSPA1L | CDH1 | CASP8 | BARD1 | NOS3 | NEUROD1 | MYH7 | CFTR |
| SKP1 | DPP4 | CCNA1 | SNCA | DLEC1 | HMMR | PLAU | IRS1 | MYLK2 | TGFB1 |
| NCL | KL | BUB1 | DRD2 | RASSF1 | NQO2 | A2M | PPARG | LMNA |  |
| YWHAB | NOX4 | NEK2 | ITGB3 | PIK3CA | BRCA2 | APP | SLC2A2 | SCN5A |  |
| HNRNPA1 | DEFB1 | INSL3 | CD44 |  |  |  |  |  |  |
| EEF1A1 | PAX8 | NDC80 | HLA-C |  |  |  |  |  |  |
| ACTG1 | CYP27B1 | RLN2 | MET |  |  |  |  |  |  |
| PABPC1 | AQP2 | PRAME | BMP2 |  |  |  |  |  |  |
| HNRNPA2B1 | HOXA9 | IL13RA2 | FGFR2 |  |  |  |  |  |  |
| RPL36AL | FXYD4 | TULP2 | EXPH5 |  |  |  |  |  |  |
| UQCRQ | UGT2A3 | HRASLS | LAD1 |  |  |  |  |  |  |
| C11orf58 (SMAP) | SLC23A3 | C19orf36 (IZUMO4) | PLEKHM1 |  |  |  |  |  |  |
| LAPTM4A | C14orf105 | GK2 | STX10 |  |  |  |  |  |  |
| ATP5G3 | SLC5A12 | CCIN | SNN |  |  |  |  |  |  |
| NACA | PRODH2 | ZNRF4 | KIAA0562 (CEP104) | |  |  |  |  |  |
| ARF1 | LRRC19 | GAGE3 | ALDH3B2 |  |  |  |  |  |  |
| B2M | UNC5CL | UBQLN3 | MPZL2 |  |  |  |  |  |  |
| PARK7 | SLC4A9 | ADAM21 | NENF |  |  |  |  |  |  |
| UQCR (UQCR11) | LOC389332 | OR7E156P | PI15 |  |  |  |  |  |  |

Ubiq: Ubiquitous, Kid: Kidney, Tes: Testis, Uter: Uterus, LC: Lung cancer, BC: Breast cancer, Alzh: Alzheimer’s, Diab: Diabetes, CM: Cardiomyopathy, CF: Cystic fibrosis.

**Coverage of protein interactions across various databases for different gene-sets**

S3 Table. Coverage of ‘total’ protein interactions for all gene-sets

|  | **Ubiq** | **Kid** | **Tes** | **Uter** | **LC** | **BC** | **Alzh** | **Diab** | **CM** | **CF** | **Average** |
| --- | --- | --- | --- | --- | --- | --- | --- | --- | --- | --- | --- |
| **STRING-API** | **44.94** | **41.04** | **42.32** | **54.02** | **44.97** | **44.23** | **42.29** | **51.94** | **38.54** | **58.84** | **46.31** |
| **STRING-Web** | 16.23 | 32.51 | 20.98 | 19.27 | 18.87 | 20.47 | 16.92 | 19.75 | 19.52 | 16.43 | **20.09** |
| **UniHI** | 12.72 | 4.28 | 5.36 | 7.51 | 7.18 | 6.46 | 18.90 | 4.92 | 5.11 | 6.44 | **7.89** |
| **iRefWeb** | **21.46** | 1.13 | 5.27 | 5.51 | 3.62 | 5.03 | 15.26 | 2.99 | **11.79** | 3.32 | **7.54** |
| **mentha** | 5.38 | 1.01 | 2.57 | 5.87 | 3.40 | 4.10 | 15.48 | 1.94 | 2.80 | 3.82 | **4.64** |
| **APID** | 5.77 | 1.18 | 3.24 | 5.14 | 3.85 | 4.15 | 16.13 | 2.41 | 3.20 | 4.00 | **4.91** |
| **HIPPIE** | 6.43 | 1.40 | 2.80 | 5.83 | 4.22 | 4.19 | 16.46 | 2.48 | 3.26 | 4.08 | **5.12** |
| **GPS-Prot** | 5.85 | 1.36 | 2.72 | 2.92 | 4.17 | 4.11 | 1.70 | 2.44 | 3.12 | 4.04 | **3.24** |
| **BioGRID** | 5.41 | 0.89 | 3.03 | 5.46 | 3.04 | 4.05 | 15.45 | 1.85 | 2.69 | 3.09 | **4.49** |
| **HPRD** | 1.65 | 0.40 | 0.45 | 1.39 | 1.39 | 0.66 | 1.14 | 0.93 | 1.72 | 0.82 | **1.06** |
| **HitPredict** | 2.92 | 0.90 | 1.92 | 3.39 | 2.21 | 1.36 | 14.62 | 1.92 | 0.78 | 3.36 | **3.34** |
| **IntAct** | 2.32 | 0.48 | 1.51 | 2.36 | 1.99 | 0.97 | 1.48 | 0.42 | 0.75 | 2.27 | **1.45** |
| **MINT** | 0.53 | 0.12 | 0.45 | 0.37 | 0.84 | 0.23 | 0.45 | 0.09 | 0.17 | 0.55 | **0.38** |
| **HINT** | 5.49 | 0.62 | 2.65 | 4.16 | 3.45 | 3.97 | 15.70 | 2.07 | 2.85 | 4.12 | **4.51** |
| **hPRINT** | **53.03** | **69.00** | **59.22** | **58.08** | **69.79** | **61.41** | **46.20** | **66.52** | **61.45** | **65.23** | **60.99** |
| **HuRI** | 0.11 | 0.54 | 0.29 | 0.01 | 0.19 | 0.22 | 0.03 | 0.04 | 0.10 | 0.25 | **0.18** |
| **IID** | 16.73 | **5.80** | **11.73** | **13.60** | **16.32** | **12.62** | **26.90** | **8.21** | 8.61 | **16.23** | **13.68** |

Ubiq: Ubiquitous, Kid: Kidney, Tes: Testis, Uter: Uterus, LC: Lung cancer, BC: Breast cancer, Alzh: Alzheimer’s, Diab: Diabetes, CM: Cardiomyopathy, CF: Cystic fibrosis. Note: Top 3 databases based on the coverage are highlighted in bold. Numbers represent percentage coverage.

S4 Table. Coverage of ‘experimentally verified’ protein interactions for all gene-sets

|  | **Ubiq** | **Kid** | **Tes** | **Uter** | **LC** | **BC** | **Alzh** | **Diab** | **CM** | **CF** | **Average** |
| --- | --- | --- | --- | --- | --- | --- | --- | --- | --- | --- | --- |
| **STRING-API** | **54.53** | **57.57** | **60.59** | **58.89** | **67.42** | **74.97** | 15.74 | **59.22** | **55.90** | **63.94** | **56.88** |
| **STRING-Web** | 25.04 | 50.14 | 39.61 | 30.40 | 36.44 | 36.27 | 15.74 | 59.22 | 37.14 | 58.08 | **38.81** |
| **UniHI** | **20.46** | **31.60** | **20.64** | **28.92** | **37.56** | **20.65** | **57.15** | **39.69** | **15.06** | **40.42** | **31.21** |
| **iRefWeb** | **47.05** | 10.02 | **23.96** | 22.94 | 19.42 | **16.71** | 47.15 | **28.64** | **39.91** | 22.76 | **27.86** |
| **mentha** | 12.09 | 9.25 | 11.16 | **24.55** | 18.48 | 13.88 | 47.88 | 19.07 | 9.47 | 26.22 | **19.20** |
| **APID** | 12.96 | 10.79 | 14.07 | 21.50 | 20.92 | 14.05 | **49.91** | 23.74 | 10.84 | 27.42 | **20.62** |
| **HIPPIE** | 14.44 | **12.83** | 12.16 | 24.37 | **22.94** | 14.18 | **50.92** | 24.44 | 11.03 | 28.02 | **21.53** |
| **GPS-Prot** | 13.14 | 12.47 | 11.82 | 12.22 | 22.66 | 13.91 | 5.27 | 24.05 | 10.55 | 27.72 | **15.38** |
| **BioGRID** | 12.15 | 8.16 | 13.14 | 22.82 | 16.50 | 13.71 | 47.78 | 18.21 | 9.10 | 21.19 | **18.28** |
| **HPRD** | 3.70 | 3.67 | 1.96 | 5.83 | 7.57 | 2.22 | 3.52 | 9.18 | 5.82 | 5.63 | **4.91** |
| **HitPredict** | 6.55 | 8.20 | 8.36 | 14.19 | 11.99 | 4.61 | 45.23 | 18.91 | 2.66 | 23.07 | **14.38** |
| **IntAct** | 5.20 | 4.35 | 6.55 | 9.87 | 10.81 | 3.27 | 4.59 | 4.12 | 2.55 | 15.55 | **6.69** |
| **MINT** | 1.20 | 1.13 | 1.94 | 1.55 | 4.56 | 0.77 | 1.39 | 0.93 | 0.59 | 3.76 | **1.78** |
| **HINT** | 12.33 | 5.71 | 11.48 | 17.39 | 18.76 | 13.44 | 48.56 | 20.39 | 9.66 | 28.25 | **18.60** |
| **hPRINT** | 6.13 | 7.34 | 2.76 | 10.67 | 11.99 | 2.96 | 4.07 | 9.65 | 2.33 | 6.16 | **6.41** |
| **HuRI** | 0.26 | 4.94 | 1.26 | 0.04 | 1.03 | 0.74 | 0.11 | 0.39 | 0.32 | 1.73 | **1.08** |
| **IID** | 13.93 | 12.51 | 12.44 | 21.37 | 22.71 | 14.38 | 48.92 | 23.81 | 10.17 | **30.13** | **21.04** |

Ubiq: Ubiquitous, Kid: Kidney, Tes: Testis, Uter: Uterus, LC: Lung cancer, BC: Breast cancer, Alzh: Alzheimer’s, Diab: Diabetes, CM: Cardiomyopathy, CF: Cystic fibrosis. Note: Top 3 databases based on the coverage are highlighted in bold. Numbers represent percentage coverage.

S5 Table. Coverage of ‘total’ protein interactions for well and less studied gene-sets

|  | **Well studied genes** | | | | | **Less studied genes** | | | | |
| --- | --- | --- | --- | --- | --- | --- | --- | --- | --- | --- |
|  | **Ubiquitous** | **Kidney** | **Testis** | **Uterus** | **Average** | **Ubiquitous** | **Kidney** | **Testis** | **Uterus** | **Average** |
| **STRING-API** | **46.38** | **37.19** | **43.88** | **56.82** | **46.07** | **41.44** | **58.63** | **33.42** | **40.26** | **43.44** |
| **STRING-Web** | 12.94 | 28.33 | 18.85 | 15.91 | **19.01** | 24.24 | 51.57 | 33.17 | 35.78 | **36.19** |
| **UniHI** | 15.35 | 4.82 | 6.02 | 8.39 | **8.64** | 6.33 | **1.82** | 1.61 | 3.19 | **3.24** |
| **iRefWeb** | **25.12** | 1.16 | 6.04 | 6.07 | **9.60** | **12.53** | 1.02 | 0.87 | 2.76 | **4.29** |
| **mentha** | 6.62 | 1.07 | 2.88 | 6.56 | **4.28** | 2.38 | 0.74 | 0.84 | 2.49 | **1.61** |
| **APID** | 6.86 | 1.19 | 3.01 | 5.65 | **4.18** | 3.14 | 1.13 | 4.59 | 2.65 | **2.88** |
| **HIPPIE** | 7.90 | 1.57 | 3.16 | 6.52 | **4.79** | 2.87 | 0.63 | 0.74 | 2.44 | **1.67** |
| **GPS-Prot** | 7.07 | 1.52 | 3.08 | 3.02 | **3.68** | 2.84 | 0.63 | 0.68 | 2.43 | **1.65** |
| **BioGRID** | 6.47 | 0.83 | 2.77 | 6.00 | **4.02** | 2.83 | 1.16 | 4.49 | 2.82 | **2.83** |
| **HPRD** | 2.10 | 0.48 | 0.51 | 1.65 | **1.18** | 0.55 | 0.06 | 0.12 | 0.13 | **0.21** |
| **HitPredict** | 3.61 | 0.97 | 2.14 | 3.77 | **2.62** | 1.23 | 0.55 | 0.68 | 1.56 | **1.01** |
| **IntAct** | 2.91 | 0.45 | 1.68 | 2.54 | **1.89** | 0.88 | 0.61 | 0.53 | 1.50 | **0.88** |
| **MINT** | 0.67 | 0.15 | 0.53 | 0.43 | **0.45** | 0.20 | 0.00 | 0.00 | 0.06 | **0.06** |
| **HINT** | 6.72 | 0.62 | 3.00 | 4.51 | **3.71** | 2.51 | 0.63 | 0.62 | 2.44 | **1.55** |
| **hPRINT** | **50.05** | **75.23** | **58.89** | **57.95** | **60.53** | **60.27** | **40.57** | **61.10** | **58.74** | **55.17** |
| **HuRI** | 0.14 | 0.38 | 0.27 | 0.00 | **0.20** | 0.04 | 1.27 | 0.43 | 0.06 | **0.45** |
| **IID** | 19.42 | **6.73** | **12.50** | **15.12** | **13.44** | 10.18 | 1.54 | **7.38** | **6.10** | **6.30** |

Note: Top 3 databases based on the coverage are highlighted in bold and underlined. Numbers represent percentage coverage.

S6 Table. Coverage of ‘experimentally verified’ protein interactions for well and less studied genes

|  | **Well studied genes** | | | | | **Less studied genes** | | | | |
| --- | --- | --- | --- | --- | --- | --- | --- | --- | --- | --- |
|  | **Ubiquitous** | **Kidney** | **Testis** | **Uterus** | **Average** | **Ubiquitous** | **Kidney** | **Testis** | **Uterus** | **Average** |
| **STRING-API** | **54.86** | **40.16** | **62.23** | **59.62** | **54.22** | **53.15** | **80.55** | **28.69** | **51.22** | **53.40** |
| **STRING-Web** | 23.31 | 40.16 | 40.17 | 28.41 | **33.01** | 32.42 | 63.30 | 28.69 | 51.22 | **43.91** |
| **UniHI** | **21.04** | **52.67** | **21.00** | **29.50** | **31.05** | **17.95** | 3.79 | 13.52 | **22.79** | **14.51** |
| **iRefWeb** | **48.33** | 14.66 | **24.82** | 23.00 | **27.70** | **41.60** | 3.89 | 7.38 | 22.28 | **18.79** |
| **mentha** | 12.99 | 14.10 | 11.16 | **24.96** | **15.80** | 8.22 | 2.84 | 11.07 | 20.23 | **10.59** |
| **APID** | 13.46 | 15.70 | 11.67 | 21.50 | **15.58** | 10.84 | 4.31 | **60.66** | 21.51 | **24.33** |
| **HIPPIE** | 15.50 | **20.72** | 12.28 | 24.80 | **18.32** | 9.90 | 2.42 | 9.84 | 19.85 | **10.50** |
| **GPS-Prot** | 13.88 | 20.08 | 11.97 | 11.50 | **14.36** | 9.81 | 2.42 | 9.02 | 19.72 | **10.24** |
| **BioGRID** | 12.70 | 11.00 | 10.76 | 22.81 | **14.32** | 9.77 | 4.42 | **59.43** | **22.92** | **24.13** |
| **HPRD** | 4.12 | 6.29 | 1.98 | 6.29 | **4.67** | 1.91 | 0.21 | 1.64 | 1.02 | **1.20** |
| **HitPredict** | 7.09 | 12.83 | 8.32 | 14.33 | **10.64** | 4.26 | 2.10 | 9.02 | 12.68 | **7.02** |
| **IntAct** | 5.71 | 5.90 | 6.53 | 9.65 | **6.94** | 3.05 | 2.31 | 6.97 | 12.16 | **6.12** |
| **MINT** | 1.32 | 1.99 | 2.04 | 1.65 | **1.75** | 0.67 | 0.00 | 0.00 | 0.51 | **0.30** |
| **HINT** | 13.18 | 8.21 | 11.65 | 17.16 | **12.55** | 8.68 | 2.42 | 8.20 | 19.85 | **9.79** |
| **hPRINT** | 7.06 | 7.73 | 2.72 | 11.05 | **7.14** | 2.18 | **6.83** | 3.69 | 6.66 | **4.84** |
| **HuRI** | 0.28 | 5.02 | 1.03 | 0.00 | **1.58** | 0.13 | **4.84** | 5.74 | 0.51 | **2.80** |
| **IID** | 14.93 | 20.00 | 12.60 | 21.53 | **17.26** | 9.68 | 2.63 | 9.43 | 19.72 | **10.36** |

Note: Top 3 databases based on the coverage are highlighted in bold and underlined. Numbers represent percentage coverage.

S7 Table. Coverage of ‘total’ exclusive protein interactions for all gene-sets

|  | **Ubiq** | **Kid** | **Tes** | **Uter** | **LC** | **BC** | **Alzh** | **Diab** | **CM** | **CF** | **Average** |
| --- | --- | --- | --- | --- | --- | --- | --- | --- | --- | --- | --- |
| **STRING-API** | **23.98** | **26.10** | **28.19** | **29.95** | **19.72** | **27.95** | **20.59** | **27.59** | **23.99** | **25.85** | **25.39** |
| **UniHI** | 1.60 | 1.20 | 1.30 | 0.65 | 0.36 | 0.82 | 0.52 | 0.88 | 0.46 | 0.85 | **0.86** |
| **iRefWeb** | **9.55** | 0.19 | 2.18 | 1.15 | 0.40 | 1.23 | 0.52 | 0.92 | **6.67** | 0.02 | **2.28** |
| **mentha** | 0.03 | 0.01 | 0.04 | 0.06 | 0.04 | 0.02 | 0.39 | 0.01 | 0.03 | 0.02 | **0.06** |
| **APID** | 0.03 | 0.02 | 0.16 | 0.08 | 0.01 | 0.03 | 0.02 | 0.03 | 0.05 | 0.01 | **0.04** |
| **HIPPIE** | 0.02 | 0.00 | 0.03 | 0.05 | 0.02 | 0.17 | 0.25 | 0.00 | 0.07 | 0.00 | **0.06** |
| **GPS-Prot** | 0.03 | 0.00 | 0.00 | 0.01 | 0.00 | 0.01 | 0.00 | 0.00 | 0.00 | 0.00 | **0.01** |
| **BioGRID** | 0.06 | 0.07 | 0.15 | 0.11 | 0.01 | 0.04 | 0.10 | 0.02 | 0.04 | 0.01 | **0.06** |
| **HPRD** | 0.06 | 0.00 | 0.00 | 0.01 | 0.01 | 0.00 | 0.00 | 0.00 | 0.01 | 0.00 | **0.01** |
| **HitPredict** | 0.06 | 0.00 | 0.02 | 0.11 | 0.01 | 0.00 | **7.19** | 0.01 | 0.00 | 0.00 | **0.74** |
| **IntAct** | 0.08 | 0.00 | 0.00 | 0.01 | 0.00 | 0.00 | 0.00 | 0.00 | 0.00 | 0.00 | **0.01** |
| **MINT** | 0.00 | 0.00 | 0.00 | 0.00 | 0.00 | 0.00 | 0.00 | 0.00 | 0.00 | 0.00 | **0.00** |
| **HINT** | 0.08 | 0.00 | 0.05 | 0.03 | 0.01 | 0.00 | 0.01 | 0.01 | 0.00 | 0.03 | **0.02** |
| **hPRINT** | **32.34** | **52.29** | **45.57** | **34.55** | **43.64** | **44.84** | **25.91** | **42.06** | **45.88** | **31.57** | **39.86** |
| **HuRI** | 0.01 | 0.32 | 0.03 | 0.01 | 0.05 | 0.01 | 0.00 | 0.00 | 0.01 | 0.01 | **0.05** |
| **IID** | 2.66 | **1.75** | **3.15** | **3.85** | **6.23** | **3.76** | 6.85 | **2.27** | 2.76 | **4.63** | **3.79** |

Ubiq: Ubiquitous, Kid: Kidney, Tes: Testis, Uter: Uterus, LC: Lung cancer, BC: Breast cancer, Alzh: Alzheimer’s, Diab: Diabetes, CM: Cardiomyopathy, CF: Cystic fibrosis. Note: Top 3 databases based on the coverage are highlighted in bold and underlined. Numbers represent percentage coverage.

S8 Table. Coverage of ‘experimentally verified’ exclusive protein interactions for all gene-sets

|  | **Ubiq** | **Kid** | **Tes** | **Uter** | **LC** | **BC** | **Alzh** | **Diab** | **CM** | **CF** | **Average** |
| --- | --- | --- | --- | --- | --- | --- | --- | --- | --- | --- | --- |
| **STRING-API** | **39.93** | **50.95** | **51.45** | **44.29** | **48.69** | **68.10** | **5.54** | **39.77** | **49.22** | **44.85** | **44.28** |
| **UniHI** | **3.76** | **20.35** | **9.56** | **5.86** | **12.83** | **6.33** | **8.72** | **14.71** | **1.34** | **14.50** | **9.80** |
| **iRefWeb** | **30.49** | 1.72 | **13.93** | **6.09** | **2.89** | **5.05** | 1.75 | **12.76** | **29.87** | 0.38 | **10.49** |
| **mentha** | 0.07 | 0.05 | 0.18 | 0.23 | 0.23 | 0.07 | 1.20 | 0.08 | 0.11 | 0.23 | **0.24** |
| **APID** | 0.09 | 0.23 | 0.56 | 0.40 | 0.09 | 0.10 | 0.08 | 0.31 | 0.16 | 0.08 | **0.21** |
| **HIPPIE** | 0.08 | 0.00 | 0.18 | 0.29 | 0.09 | 0.57 | 0.95 | 0.00 | 0.24 | 0.00 | **0.24** |
| **GPS-Prot** | 0.08 | 0.05 | 0.00 | 0.04 | 0.05 | 0.07 | 0.00 | 0.00 | 0.00 | 0.00 | **0.03** |
| **BioGRID** | 0.17 | 0.77 | 0.56 | 0.51 | 0.05 | 0.17 | 0.29 | 0.23 | 0.13 | 0.08 | **0.30** |
| **HPRD** | 0.13 | 0.00 | 0.00 | 0.03 | 0.05 | 0.00 | 0.00 | 0.00 | 0.03 | 0.00 | **0.02** |
| **HitPredict** | 0.22 | 0.00 | 0.12 | 0.45 | 0.05 | 0.00 | **22.29** | 0.08 | 0.00 | 0.00 | **2.32** |
| **IntAct** | 0.19 | 0.00 | 0.00 | 0.04 | 0.00 | 0.00 | 0.00 | 0.00 | 0.03 | 0.00 | **0.03** |
| **MINT** | 0.00 | 0.00 | 0.00 | 0.00 | 0.00 | 0.00 | 0.00 | 0.00 | 0.00 | 0.00 | **0.00** |
| **HINT** | 0.19 | 0.05 | 0.28 | 0.46 | 0.23 | 0.03 | 0.11 | 0.31 | 0.03 | 0.68 | **0.24** |
| **hPRINT** | 0.42 | **3.22** | 0.28 | 3.86 | 0.70 | 0.10 | 0.53 | 0.70 | 0.16 | 0.68 | **1.06** |
| **HuRI** | 0.03 | 3.04 | 0.14 | 0.02 | 0.28 | 0.03 | 0.02 | 0.00 | 0.03 | 0.23 | **0.38** |
| **IID** | 0.26 | 2.18 | 1.60 | 2.59 | 2.52 | 1.92 | 3.94 | 2.65 | 0.70 | **4.28** | **2.26** |

Ubiq: Ubiquitous, Kid: Kidney, Tes: Testis, Uter: Uterus, LC: Lung cancer, BC: Breast cancer, Alzh: Alzheimer’s, Diab: Diabetes, CM: Cardiomyopathy, CF: Cystic fibrosis. Note: Top 3 databases based on the coverage are highlighted in bold and underlined. Numbers represent percentage coverage.

S9 Table. Coverage of exclusive protein interactions using STRING-Web for selected gene-sets

| **Database** | **Total interactions** | | | | | | **Experimental interactions** | | | | | |
| --- | --- | --- | --- | --- | --- | --- | --- | --- | --- | --- | --- | --- |
|  | **Ubiq** | **Kid** | **BC** | **Alzh** | **CM** | **Average** | **Ubiq** | **Kid** | **BC** | **Alzh** | **CM** | **Average** |
| **STRING-Web** | **5.66** | **19.57** | **9.41** | 4.88 | **8.81** | **9.67** | **13.06** | **43.52** | **22.47** | **5.54** | **29.41** | **22.80** |
| **UniHI** | 1.87 | 1.21 | 0.86 | 0.61 | 0.49 | **1.01** | **4.37** | **20.35** | **6.33** | **8.72** | **1.34** | **8.22** |
| **iRefWeb** | **10.41** | 0.19 | 1.26 | 0.52 | **6.94** | **3.87** | **31.88** | 1.72 | **5.05** | 1.75 | **30.03** | **14.09** |
| **mentha** | 0.03 | 0.01 | 0.02 | 0.40 | 0.03 | **0.10** | 0.07 | 0.05 | 0.07 | 1.20 | 0.11 | **0.30** |
| **APID** | 0.03 | 0.02 | 0.03 | 0.02 | 0.05 | **0.03** | 0.09 | 0.23 | 0.10 | 0.08 | 0.16 | **0.13** |
| **HIPPIE** | 0.02 | 0.00 | 0.17 | 0.30 | 0.07 | **0.11** | 0.08 | 0.00 | 0.57 | 0.95 | 0.24 | **0.37** |
| **GPS-Prot** | 0.03 | 0.005 | 0.01 | 0.000 | 0.000 | **0.01** | 0.08 | 0.05 | 0.07 | 0.00 | 0.00 | **0.04** |
| **BioGRID** | 0.07 | 0.07 | 0.04 | 0.10 | 0.04 | **0.06** | 0.17 | 0.77 | 0.17 | 0.29 | 0.13 | **0.31** |
| **HPRD** | 0.06 | 0.00 | 0.00 | 0.00 | 0.01 | **0.01** | 0.13 | 0.00 | 0.00 | 0.00 | 0.03 | **0.03** |
| **HitPredict** | 0.07 | 0.00 | 0.00 | **7.20** | 0.00 | **1.45** | 0.25 | 0.00 | 0.00 | **22.29** | 0.00 | **4.51** |
| **IntAct** | 0.08 | 0.00 | 0.00 | 0.00 | 0.00 | **0.02** | 0.19 | 0.00 | 0.00 | 0.00 | 0.03 | **0.04** |
| **MINT** | 0.00 | 0.00 | 0.00 | 0.00 | 0.00 | **0.00** | 0.00 | 0.00 | 0.00 | 0.00 | 0.00 | **0.001** |
| **HINT** | 0.08 | 0.00 | 0.00 | 0.01 | 0.00 | **0.02** | 0.19 | 0.05 | 0.03 | 0.11 | 0.03 | **0.08** |
| **hPRINT** | **38.16** | **54.74** | **49.40** | **32.09** | **49.17** | **44.71** | 0.46 | **3.22** | 0.10 | 0.53 | 0.16 | **0.89** |
| **CCSB** | 0.01 | 0.32 | 0.01 | 0.00 | 0.01 | **0.07** | 0.03 | 3.04 | 0.03 | 0.02 | 0.03 | **0.63** |
| **IID** | 3.17 | **1.82** | **4.25** | **7.61** | 2.96 | **3.96** | 0.28 | 2.18 | 1.92 | 3.94 | 0.70 | **1.80** |

Ubiq: Ubiquitous, Kid: Kidney, BC: Breast cancer, Alzh: Alzheimer’s, CM: Cardiomyopathy. Note: Top 3 databases based on the coverage are highlighted in bold and underlined. Numbers represent percentage coverage.
